# Charting the transition from in vitro gliogenesis to the in vivo maturation of transplanted human glial progenitor cells

**DOI:** 10.1101/2025.04.22.649367

**Authors:** John N. Mariani, Steven J. Schanz, Benjamin Mansky, Xiaolu Wei, Carter C. Long, Devin Chandler-Militello, Nguyen P.T. Huynh, Steven A. Goldman

## Abstract

Neither rodent models nor in vitro studies of human cells adequately describe the molecular ontogeny of human glial progenitor cells (hGPCs). Here, we used scRNA-seq together with scATAC-Seq and CUT&TAG assessment of chromatin availability to track the in vitro genesis and in vivo differentiation of hGPCs from pluripotent stem cells (PSCs). In vitro, the hGPC pool comprised 4 transcriptionally-distinct subpopulations, each associated with a distinct pattern of chromatin accessibility and histone modification of stage-dependent genes. After the neonatal transplant of these cells into myelin-deficient shiverer mice, they differentiated further as astrocytes and oligodendrocytes. A combination of gene co-expression, motif enrichment, cell-trajectory, and cell-cell interaction analyses revealed that the host environment potentiated the context-dependent differentiation of the hGPCs, via their activation of distinct gene regulatory networks. Together, these data chart the process by which human PSC-derived GPCs are generated in vitro and diversify in vivo to mature as astrocytes and oligodendrocytes.

## INTRODUCTION

The loss of functional astrocytes and oligodendrocytes contributes to a broad variety of neurological and psychiatric disorders. Both in development and beyond, these glial cells can be generated from bipotential oligodendrocyte-astrocyte glial progenitor cells (GPCs), a pool of which remains in the adult human brain parenchyma, but which may be either depleted or dysfunctional in the setting of disease^1–6^. As a result, human GPCs (hGPCs) have emerged as attractive cellular therapeutic candidates for the treatment of glial-based disease ^7,8^. To explore the potential utility of these cells as transplant vectors, we previously established a human glial chimeric mouse model by which to study both the behavior of disease-derived human glia in vivo ^9–11^, and their potential replacement by healthy donor hGPCs. While these studies have established the capacity of transplanted hGPCs to repopulate and myelinate host brain ^9,12,13^, the transcriptional and epigenetic responses of these cells to being transplanted have not been well-defined, nor have the changes in population composition between cultured and engrafted hGPCs been deeply explored.

In this study, we leveraged single cell RNA sequencing (scRNA-seq) and bulk cleavage under targets and tagmentation (CUT&Tag) sequencing to define both the cell composition and transcriptional regulation of hGPCs at various stages of glial differentiation and maturation. Starting with two cell sources - one embryonic stem cell line (WA09), and one induced pluripotent stem cell line (C27) - we generated hGPCs using our established induction protocol ^11^. Using scRNA-Seq in tandem with scATAC-Seq analysis of chromatin accessibility and CUT&TAG assessment of histone modifications, we assessed both the transcriptional and epigenetic distinctions among hGPCs produced in vitro, and identified four readily distinguishable stages of hGPC ontogeny, downstream of multilineage-competent neural progenitor cells. We then transplanted these cells into the corpora callosa of neonatal shiverer mice, and sacrificed the mice 19 weeks later, isolating both human and mouse cells from the engrafted forebrains. At that point, we assessed the transcriptional distinctions between cultured hGPCs and those extracted back from the murine brain after long-term residence in vivo, and used these data to identify those context-dependent gene regulatory networks that directed the diversification and terminal glial differentiation of hGPCs in vivo. By this means, we have established an epigenetic and transcriptional roadmap by which to describe the ontogeny of human glia from pluripotent stem cells, while highlighting the importance of tissue-derived, contextual cues in both potentiating and directing terminal astrocytic and oligodendrocytic maturation in vivo.

## RESULTS

### In vitro GPC differentiation yields compositions free of pluripotent signatures

We first used scRNA-seq to profile the differentiation of hGPCs from pluripotent stem cells (PSCs), including both embryonic stem cells (hESCs; line WA09) and induced PSCs (iPSCs; line C27), using scRNA-seq (10X Genomics, v3.1) (**Fig. 1A**). Using our described in vitro differentiation protocol, which entails a 6-stage, 130-160 day procedure by which to generate hGPCs^11^, we first confirmed that gene expression patterns of human PSCs and hGPCs were wholly distinct, with no residual contributions of the starting PSC lines to the expression signatures of their derived glia (**Figs. 1B-D**). Canonical markers of pluripotency, LIN28A and POU5F1 (also known as OCT4), were highly expressed in PSCs but were effectively undetectable at the hGPC stage (**Fig. 1E**)^14,15^. Indeed, co-expression of LIN28A and POU5F1 was detected in 99.4% of PSCs (17,105 PSCs), whereas not a single hGPC among 37,805 cultured GPC-stage cells co-expressed these markers (**Fig. 1F**). The eradication of PSCs was likely the product of both the strong pro-differentiative signals imparted by our induction protocol, and the relatively extended period of time during which these cells were allowed to mature as hGPCs in vitro before their harvest and assessment.

**Figure 1.**
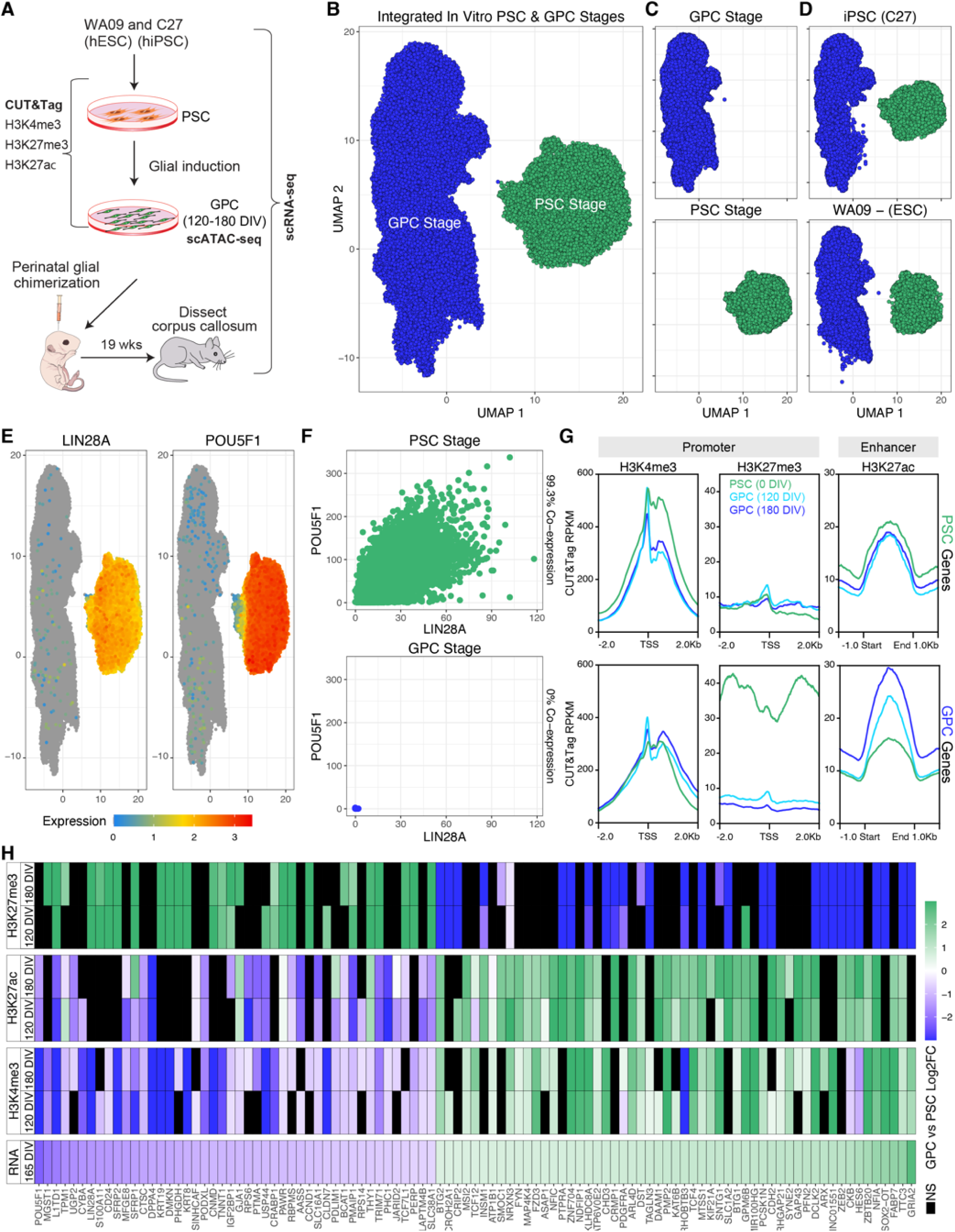
hGPC differentiation eliminates pluripotent signatures, and is orchestrated epigenetically. **A.** Human induced pluripotent stem cells (iPSCs; line C27) and embryonic stem cells (ESCs; line WA09) were captured either as naïve pluripotent stem cells, or after differentiation into glial progenitor cells (GPCs; 120-180 days in vitro). Both were then assessed by scRNA-Seq, and by bulk Cleavage Under Targets & Tagmentation (CUT & Tag) for H3K4me3, H3K27me3, and H3K27ac. **B.** Dimensionality reduction by UMAP of integrated PSC and GPCs, **C)** split by stage, and **D)** split by line (n=2 PSCs, n=4 GPCs). The complete abrogation of pluripotent signatures from the differentiated hGPC products of both lines is notable. **E-F.** Expression and co-expression of the pluripotency genes LIN28A and POU5F1 is limited to PSCs, as illustrated on integrated UMAPs (**E)**, and as recorded with raw RNA counts (**F**; co-expression is noted on the right axis). **G.** Histone modification profiles at annotated regulatory regions of both PSC and two hGPC stages (indicated by line color, n=5-7 per mark per timepoint). PSC- or GPC-linked modifications were determined via differential expression (MAST; FDR<0.01, Log_2_FC>0.25). H3K4me3 and H3K27Ac generally serve as activating marks, whereas H3K27me3 is repressive. **H.** Heatmap shows those genes differentially expressed between stages; filtered for those which had at least 2 concordant CUT & Tag marks differentially bound between PSCs and GPCs. Black indicates non-significant loci.

### In vitro GPC differentiation is orchestrated epigenetically

Next, we sought to identify those transcriptional changes that evolved during hGPC differentiation in vitro. Differential expression (DE) analysis with MAST identified 956 genes enriched in PSCs and 1,189 in GPC cultures, irrespective of cell line (FDR <0.01, log_2_ fold-change >0.25)^16^. Among DE genes in PSCs were POU5F1, ESRG and LIN28A; in contrast, the GPC pool upregulated markers of glial development, such as OLIG1, PDGFRA, NFIA, and NFIB ^15,17–19^ (**Supplementary Data 1**). Given these transcriptional differences, we proceeded to examine the epigenetic landscapes of both the PSC and hGPC stages, using bulk CUT&Tag for H3K4me3, H3K27me3 and H3K27ac^20^. For hGPCs, we performed CUT&Tag at both 120 and 180 DIV, to observe those histone modifications that arose during progressive hGPC maturation (**Supplementary Data 1**). Typically, H3K4me3 is enriched at active promoters, H3K27ac marks active enhancers, and H3K27me3 acts as a repressive mark near transcription start sites (TSS).

We found that the promoters of genes selectively enriched in PSCs relative to hGPCs showed the highest average binding of H3K4me3 in PSCs, which declined with differentiation (**Fig. 1G**). Enhancers of annotated PSC-enriched genes also displayed higher binding of H3K27ac in PSCs. However, repression via H3K27me3 binding of PSC-enriched promoters appeared similar across stages, suggesting that the loss of pluripotent gene expression at the hGPC stage is linked to a reduction in active marks (H3K4me3 and H3K27ac), rather than with an increase in repressive marks. This pattern is evident in POU5F1, in which dramatic decreases in H3K4me3 and H3K27ac are observed during glial differentiation, with minimal changes in H3K27me3 (GH06J031161, **Supplementary** Figs. 1A-B). In contrast, promoters of hGPC enriched genes exhibited higher average H3K4me3 binding and progressively increasing H3K27ac enhancer binding as glial differentiation progressed (**Fig. 1G**). Furthermore, repressive H3K27me3 binding at the promoters of hGPC enriched genes fell progressively with glial differentiation, suggesting the stage-linked derepression of these loci. By way of example, the glial enriched gene NFIA exhibited greater H3K4me3 binding during differentiation, which was accompanied by a dramatic loss of H3K27me3 binding at its promoter, along with increased H3K27ac enrichment at one of its putative enhancers (GH01J060989, **Supplementary Figs. 1C-D**).

We next intersected the differentially-expressed histone modifications with those genes whose expression levels differed concordantly across stages; to do so, we focused on those differentially expressed genes that were most regulated across our three inspected histone marks (**Fig. 1H**). Of these, 934 genes exhibited differential enrichment in at least one of the histone marks (H3K4me3, 496; H3K27me3, 243; K3K27ac, 564), with 72 found to be concordantly engaged by all three. Of note, GPC-enriched genes such as GRIA2, FABP7, NFIA, NFIB, NFIC, SLC1A2, and PDGFRA were each modulated – with appropriate directionality - by at least two marks. Similarly, PSC-enriched genes like POU5F1, LIN28A, PODXL, and MGST1 were regulated by at least two of these marks. These findings underscore the profound modification of PSCs by our hGPC differentiation protocol, effectively eliminating residual PSCs in a manner that is largely orchestrated through epigenetic mechanisms.

### hGPCs are comprised of distinct stage-defined subpopulations

Both murine and human oligodendrocytes and their progenitors have been reported to exhibit significant heterogeneity in vivo ^21,22^. With that in mind, we next sought to deeply characterize the composition and heterogeneity of our PSC-derived hGPC preparations. To do so, we integrated the scRNA-Seq expression profiles of cultured hGPCs with those extracted back from adult mouse brains into which they had been neonatally engrafted. This integrated analysis allowed us to accurately define the composition of our in vitro preparations, while simultaneously assessing the degree of subsequent, context-dependent differentiation driven by the in vivo environment.

To this end, we utilized our previously-established chimeric mouse model, wherein hGPCs were transplanted perinatally into the corpus callosum of immunodeficient and myelin-deficient shiverer mice (rag2^-/-^ x MBP^shi/shi^) ^9,23–27^. We had previously shown that hGPCs transplanted into these mice reliably differentiate into both myelinating oligodendrocytes and astrocytes, with a complement of persistent hGPCs^11^. In addition, we had noted that donor cell colonization was especially robust in the callosal and capsular white matter of these mice. On that basis, we transplanted hGPCs into neonatal immunodeficient shiverers and killed them at 19-20 weeks, allowing sufficient time to permit the in vivo expansion and maturation of the engrafted hGPCs. We then extracted both human and mouse cells back from the callosal white matter of these mice for scRNA-seq analysis (n=4 for WA09, n=3 for C27). Following alignment to a dual-species reference with STARsolo^28^, human and mouse cells were then identified bioinformatically, utilizing a custom approach for downstream analysis of either species (see Methods). Leiden clustering of both integrated stages revealed four transcriptionally-distinct hGPC cell states (GPCs1-4), as well as one resembling early neural progenitor cells (NPC), and three additional clusters comprising astrocytes, immature oligodendrocytes (imOL), and mature oligodendrocytes (maOL) (**Figs. 2D-F, Supplementary Fig. 2A**)^29^. These cell states emerged similarly across both cell lines in culture (**Supplementary** Figs. 2B-C) and were apparent whether or not the cultures were pre-enriched for hGPCs by CD140a-based FACS, an exploratory step that we found depleted only a fraction of the NPCs, without substantively affecting our transcriptional data otherwise (**Supplementary Fig. 2D**).

**Figure 2.**
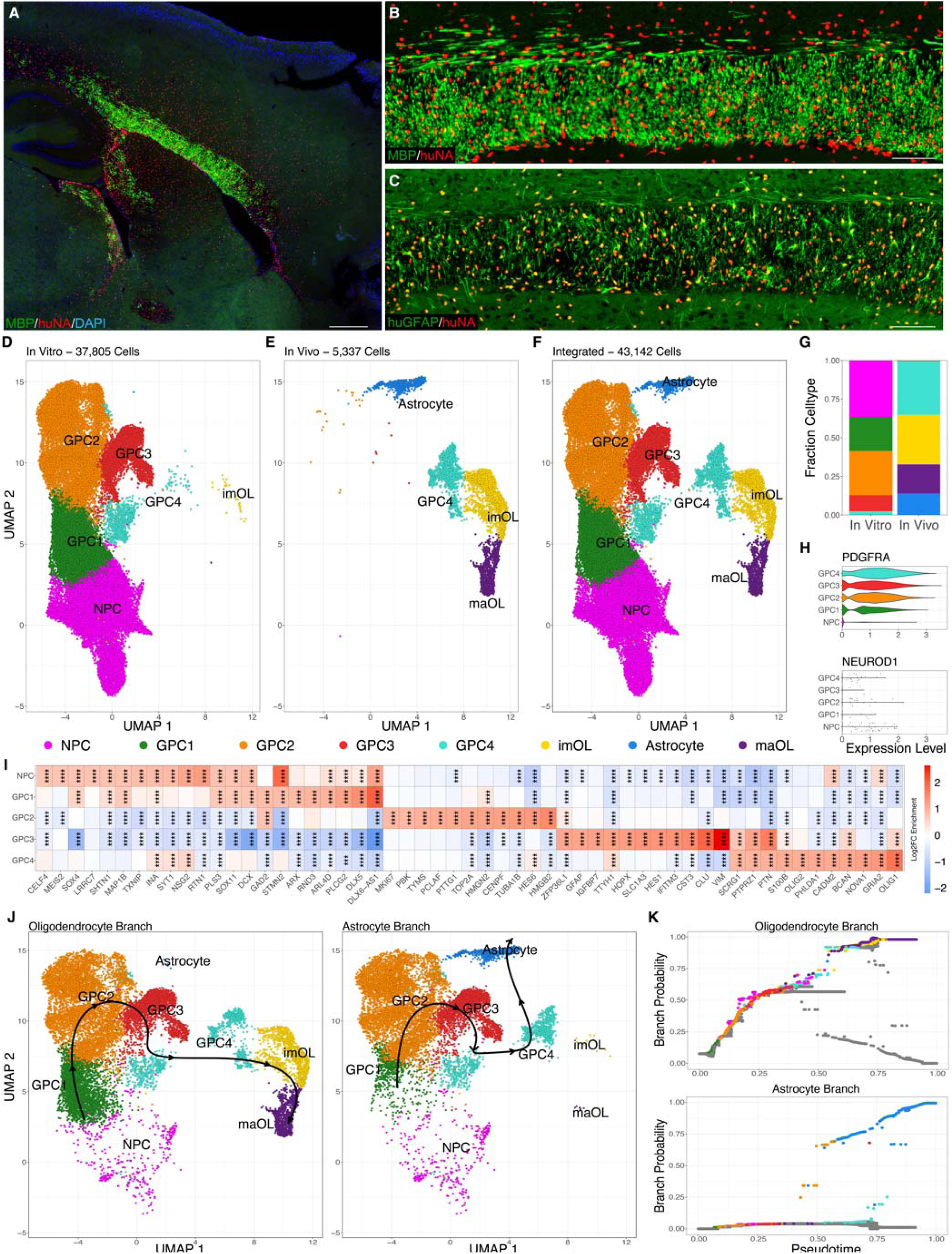
hGPCs differentiate further in vivo. After transplantation into shiverer corpus callosum, hGPCs undergo further differentiation 19 wks post-graft into MBP^+^ oligodendrocytes (**A-B**) and GFAP^+^ astrocytes (**C**). A sagittal section is shown in (**A**) and higher power callosal images in (**B-C**). **D-E.** Integrated UMAP representation of identified cells, either (**D**) in vitro, after GPC differentiation (165 DIV), or (**E**) in vivo, after neonatal transplant (n=6 in vitro GPCs, n=7 in vivo). **F.** Integrated UMAP including cells identified both in vitro and in vivo. **G.** Cell type compositions of in vitro and in vivo samples. **H.** Violin plots of PDGFRA define GPCs in vitro, while the lack of NEUROD1 expression indicates the absence of neurons in these compositions. **I.** Heatmap of top 11 enriched genes per cell type in vitro, highlighting distinctions between 4 major identified hGPC subpopulations (MAST; FDR <0.01, Log_2_FC >0.25). **J-K.** UMAPs with overlaid trajectories and branch probability plots across pseudotime of oligodendrocyte and astrocyte lineages, calculated with Palantir^33^. huNA: human nuclei, *imOLs and mOLs: immature and mature oligodendrocytes.* **** p-value < 0.0001. Scale bars: **A**, 500 µm, **B-C**, 100 µm.

We next assessed the effect of transplantation on the composition of these hGPCs, with an eye towards assessing the effect of the in vivo environment on the differentiation trajectories of each hGPC subclass. We found that whereas the hGPC cultures included all four GPC cell states as well as NPCs, along with rare immature oligodendrocytes (**Supplementary** Figs. 2D,G), the human cells extracted back from chimeras were comprised almost entirely of GPC4-classified hGPCs, along with substantial fractions of imOLs, maOLs, and astrocytes (**Figs. 2E,G**). These data revealed a pronounced shift in identity upon engraftment and expansion in the murine white matter.

This integrated overview of both cultured and in vivo hGPC expression then allowed us to define the relationship of hGPC transcriptional states in culture to their counterparts after in vivo engraftment. To that end, we first focused on the composition of our hGPC preparations at their time of harvest for transplant, at 160 DIV. We assessed DE by each of the five primary cell states, relative to the remaining culture subpopulations (MAST, FDR < 0.01, Log_2_ fold-change >0.25, **Supplementary Data 2**); by this means, we sought to identify the defining transcriptional features of each state (**Fig. 2I**; top 11 genes per cell state shown). We found that NPCs differentially expressed a set of early neuronal genes, yet they did not demonstrate significant expression of more lineage-restricted neuronal genes, such as NEUROD1 (**Fig. 2H**). Among GPC1-categorized cells, the GPC marker PDGFRA first appeared appreciably, accompanied by the decreased expression of neuronal lineage markers (**Figs. 2H-I**)^24,30^, despite the persistent expression of early, lineage-agnostic neural genes. GPC2 included the primary actively cycling population within the culture, marked by enrichment of active proliferation markers including MKI67, TOP2A, and CENPF, and by the highest G2M and S phase scores of all states (**Fig. 2I, Supplementary Figs. 2E-F**). Within GPC3, there was a notable enrichment of genes typically observed in both radial glia and astrocytes, including GFAP, SLC1A3 (Also known as GLAST), HOPX, CLU, and VIM. Lastly, GPC4 was distinguished by the most specified hGPC signature, with hGPCs from chimeric white matter primarily belonging to this cluster. In particular, expression of OLIG1, OLIG2, BCAN, PDGFRA, and PTPRZ1 were all significantly enriched in this population, a signature typifying bipotential oligodendrocyte-astrocyte hGPCs ^31^.

### Patterns of chromatin accessibility predict hGPC heterogeneity

Given the transcriptional heterogeneity of cultured hGPCs, we asked whether the major identified classes of hGPCs could be similarly predicted on the basis of their patterns of chromatin accessibility. To this end, we utilized scATAC-seq (10X Genomics) of 180 DIV ESC-derived hGPCs (WA09)^32^, where subpopulation assignment was informed by our scRNA-seq data (**Supplementary Fig. 3A**) Enrichment of differentially accessible regions of chromatin within each subpopulation was determined using Wilcoxon rank sum tests (adjusted p-value <0.01, **Supplementary Data 2**) revealing substantial concordance between regions of open chromatin and differentially expressed genes. By way of example, among the significant peaks, the GPC4 transcriptional markers NKX2-2 and OLIG2 were found to be preferentially accessible in GPC4-class cells relative to other subpopulations (**Supplementary Fig. 3B**).

We next calculated gene activity enrichment using Wilcoxon rank sum tests of each subpopulation vs. all other nuclei (Adjusted p-value < 0.01, **Supplementary Data 2**). Inspection of those genes most enriched by scATAC-Seq-inferred gene activity (**Supplementary Fig. 3C**) that were also differentially expressed (**Supplementary Fig. 3D**), revealed similar state assignments as those based on gene expression alone. These included the enrichment of neural signatures in NPCs and GPC1, proliferation-associated expression (e.g., CDK6, RPN2, RRM2, HAT1) in GPC2, and astrocyte-associated enrichment (e.g., GFAP, TNC, SOX9) in GPC3. GPC4, the most differentiated subpopulation of hGPCs in our cultures, exhibited relative accessibility at a host of GPC genes; gene expression (scRNA-seq) and predicted gene activity (scATAC-seq) in GPC4 revealed that those genes significantly enriched in both assays were almost entirely concordant (97.3%; **Supplementary Fig. 3E**, *shown in blue*). These included the GPC4-enriched genes OLIG1, OLIG2, SOX10, NKX2-2, GRIA2, NOVA1, BCAN, S100B, ADGRL3, CCND1, PCDH17, LRRC4C, THY1, and CA10. Conversely, GPC4 was also characterized by the downregulation of early astrocytic (e,g,, NFIA, NFIB, CLU) and neural (DLX1, DLX2, GAD2) genes, along with diminished chromatin accessibility. Interestingly, while upregulated transcriptionally, the mature GPC-associated genes PDGFRA, PTPRZ1, DLL3, and FYN were among those whose chromatin accessibility was not significantly regulated, suggesting their direct transcriptional regulation. In contrast, the transcription of prototypical hGPC genes including PCDH15, CSPG4 (Also known as NG2), FGF14 and GPR17 was not significantly upregulated in GPC4, even while their chromatin accessibility was enriched, suggesting that these genes may become primed in vitro for later, differentiation-associated upregulation in vivo.

### Transplanted heterogeneous populations of hGPCs undergo convergent maturation in vivo

We next employed Palantir to predict the diverse differentiation trajectories existing within our hGPC cultures, and their progress towards glial phenotype in vivo^33^. This analysis unveiled three trajectories: oligodendrocytic, astrocytic, and one toward more differentiated NPCs in vitro (**Figs. 2J-K, Supplementary** Figs. 2G-H). The trajectory leading to mature oligodendrocytes was predicted to originate from GPC1/NPC cells, transitioning through GPC2, GPC3, and culminating in GPC4 (**Fig. 2J**) before differentiating further in vivo. Indeed, a discernible gap in branch probability - a measure of how likely a cell is to differentiate into the branch’s terminal cell type (**Fig. 2K**) - was observed in GPC4 between its in vitro and vivo stages, predicting the further oligodendrocytic differentiation of these hGPCs after their in vivo residence. Accordingly, the hGPCs in this branch differentiated into both immature and mature oligodendrocytes (imOLs and maOLs) in vivo.

Remarkably, the trajectory towards astrocytic phenotype followed a comparable path, with cells transitioning from GPC4 in vitro to GPC4 in vivo, rather than directly from another in vitro population such as the more astrocytic GPC3 (**Fig. 2J**). Branch probabilities along this trajectory exhibited greater divergence compared to the oligodendrocyte branch, suggesting the emergence of transitional astroglial progenitors after transplant (**Fig. 2K**). In addition, a third minor in vitro trajectory was predicted, whereby a fraction of less mature cultured NPCs upregulated early neural genes such as SYT1, MEIS2, and BCL11B, without exhibiting evidence of further neuronal commitment (**Supplementary Fig. 2G**). Accordingly, neither donor-derived NPCs nor neurons were ever recovered from chimeric white matter in this study. Taken together, these data highlight the heterogeneity of hGPC phenotypes within these preparations, and the relatively restricted range of glial phenotypes into which these subpopulations differentiate in vivo after transplantation.

### Human donor cells can all be recovered from host brains as oligodendrocytes, astrocytes, and hGPCs

We next sought to characterize in more detail those human cells extracted from chimeric host brains. Re-integration and re-clustering of these cells identified seven cell states (**Figs. 3A, Supplementary** Figs. 4A-B) that were detected from both cell lines (**Fig. 3B**). These cell types encompassed PDGFRA-enriched hGPCs, MKI67-enriched cycling hGPCs (cGPCs) and cycling astrocyte progenitor cells (cAPC), BCAS1-enriched imOLs, MOBP-enriched mOLs, SLC1A2 (Also known as GLT1)-enriched immature astrocytes (imAstrocytes), and GFAP and AQP4-enriched mature astrocytes (**Figs. 3A-C**; **Supplementary Fig. 4C**).

**Figure 3.**
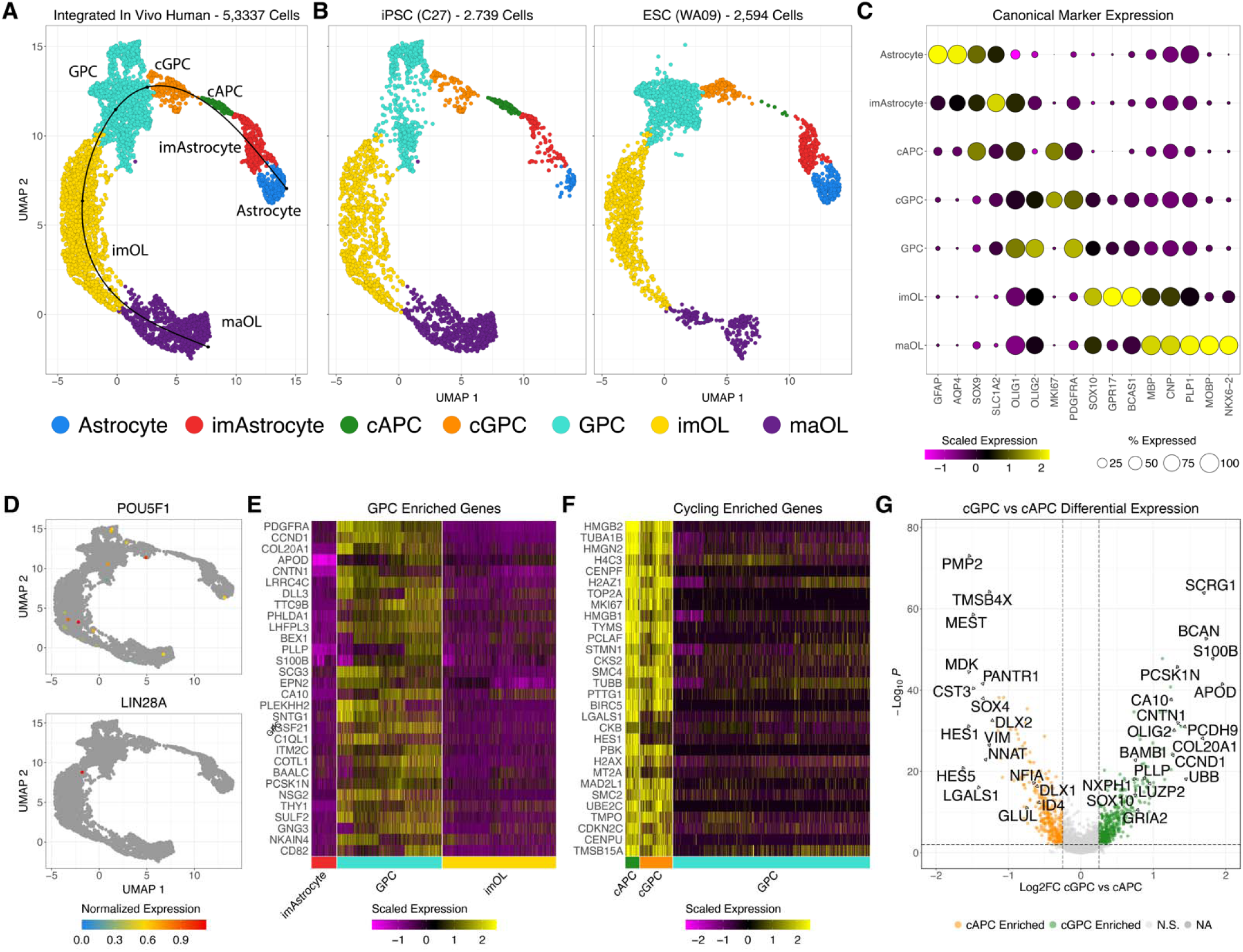
Human cells in chimeric white matter differentiate solely as GPCs, oligodendrocytes, and astrocytes. After in vivo residence, human donor cells have all differentiated as oligodendrocytes, astrocytes, or their lineage restricted progenitors. **A.** Integrated UMAP representation of identified cells extracted from chimerized shiverer brains, 19 weeks after neonatal transplant. Principal curve is pseudotime progression from slingshot. **B.** Comparable cell types identified in samples derived from ESCs or iPSCs. **C.** Dot plot depicting the scaled expression of canonical markers in identified cell types. **D.** Feature plots depicting low to no level of pluripotent marker expression with no co-expression. **E.** Top genes differentially enriched in hGPCs that are rapidly downregulated in both imOLS and imAstrocytes. **F.** Shared differentially enriched genes in proliferating (cycling) cGPCs and cAPCs vs GPCs. **G**. Volcano plot of genes differentially expressed between cGPCs and cAPCs. Differential expression was conducted with MAST (FDR < 0.01, Log_2_FC > 0.25). *cGPC: cycling GPCs; GPC: glial progenitor cells; imOL: immature oligodendrocytes; maOL: mature oligodendrocyte; cAPC: cycling astrocyte progenitor cells; imAstrocyte: immature astrocytes*.

We next used pseudotime analysis to estimate the lineage progression within and between these identified cell types, both in vitro and in vivo. This analysis focused on both cycling populations, and progressed toward their mature oligodendrocytic or astrocytic endpoints (**Fig. 3A**)^34^; it confirmed that the expression of the pluripotency markers POU5F1 and LIN28A was effectively extinguished by donor cells in vivo, with no extracted human cells co-expressing both (**Fig. 3D**; among 5,337 human cells). Thus, the perinatal transplantation of human iPSC- or ESC-derived GPCs resulted in their colonization and differentiation, with no evidence of persistent pluripotent or other off-target lineage cells appearing during roughly 5 months of in vivo residence.

### Transcriptionally-distinct cycling pools of bipotential and astrocyte-biased progenitors appear in vivo

We next focused on defining the expression profiles of hGPCs during both their mitotic expansion and subsequent quiescence in vivo. We assessed differential gene expression between hGPCs and imOLs and imAstrocytes (MAST, FDR <0.01, log_2_FC >0.25, **Supplementary Data 3**), and investigated the 193 intersecting hGPC-enriched genes that were downregulated during *both* astrocytic and oligodendrocytic differentiation (**Fig. 3E**). Human GPCs in chimeric white matter were highly enriched for a host of hGPC genes, that included PDGFRA, CCND1, APOD, CNTN1, PLLP, OLIG1, OLIG2, PTPRZ1, CA10, COL20A1, and NXPH1^35^; these cells co-clustered, and did not significantly express genes associated with active mitotic expansion. Yet besides these quiescent hGPCs, two discrete populations of cycling progenitors were also identified, which we designated as cGPCs and cAPCs. These populations showed high G2M and S phase scores relative to other human populations (**Supplementary Fig. 4D**) and were enriched for proliferation-associated genes. Overall, 268 intersecting cGPC and cAPC genes were enriched relative to quiescent GPCs (MAST, FDR <0.01, log_2_FC >0.25) (**Fig. 3F, Supplementary Data 3**).

While similar in proliferation-associated gene expression, cGPCs and cAPCs differed significantly in gene expression (**Fig. 3G**), which appeared to predict their progression toward oligodendrocytic or astrocytic fates, respectively (MAST, 502 genes enriched in cGPCs, 351 enriched in cAPCs, FDR <0.01, log_2_FC >0.25, **Supplementary Data 3**). cGPCs were enriched for early oligodendroglial transcripts that included OLIG2, BCAN, PCDH9, SOX10, APOD, BAMBI, and PLLP. In contrast, cAPCs were enriched for early astrocytic genes including SOX9, HES5, ID4, NFIA and GLUL. Together, these data suggest that in vivo, cycling hGPCs fall into transcriptionally-distinct subpopulations, whose transcriptional signatures may bias – and perhaps predict - their subsequent oligodendroglial or astroglial fate.

### Context-dependent differentiation of human GPCs in callosal white matter

Given the progressive differentiation of GPCs after transplantation, we next sought to define the transcriptional differences between the cultured and engrafted GPCs, so as to identify transcriptional activators driving context-dependent differentiation. We focused on the GPC4 population of our integrated in vitro and in vivo data, as these GPCs were the most similar in the two contexts (**Fig. 2F**; compare to **Fig. 4A**). Furthermore, GPC4 was predicted to comprise the cluster through which differentiation occurred in our cell trajectory analysis (**Figs. 2G-H**). Indeed, as previously noted, reanalysis of in vivo human cells alone found that 99.2% of engrafted GPC4 cells were identifiable as GPCs or cGPCs (**Fig. 3A**, **Supplementary Fig. 4C**). Analysis of differential expression by this population revealed 685 hGPC genes that were upregulated in the in vivo environment, whereas cultured hGPCs were enriched for 320 (MAST, FDR <0.01, Log_2_FC >0.25, **Supplementary Data 4**). The GPCs in vivo upregulated a cohort of oligodendrocytic lineage genes that included SOX10, OLIG2, OLIG1, PCDH9, BCAN, PDGFRA, CCND1, PTPRZ1, while they downregulated genes typically associated with neural identity (**Fig. 4B**), suggesting that the murine environment strongly potentiated the glial maturation of transplanted GPCs. Together, these data suggest that the extensive differences in gene expression observed in hGPCs in vivo, compared to their counterparts in vitro before transplant, were in response to the tissue environment.

**Figure 4.**
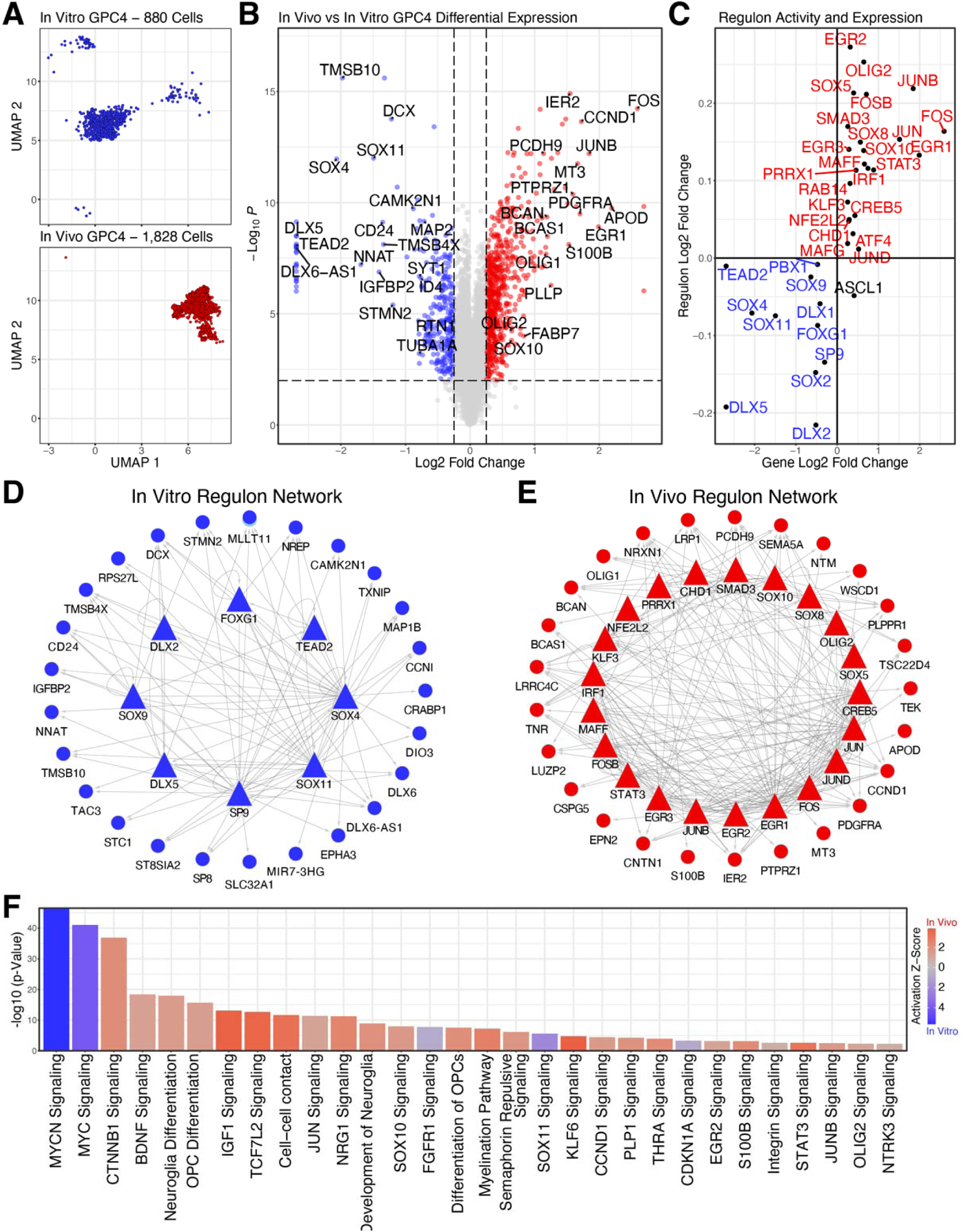
Regulation of in vivo human GPC differentiation. **A.** UMAP plots the GPC4 population of Fig. 2A, split by in vitro and in vivo, the latter in mouse callosal white matter, 19 weeks after neonatal transplant. This subpopulation comprises both the most mature hGPCs in vitro, and nearly all of the in vivo hGPCs and cGPCs. **B.** Differential expression between GPC4-subgroup hGPCs in vitro and in vivo (MAST, FDR < 0.01, Log_2_FC.> 0.25). **C.** Scatter plot of pySCENIC calculated transcription factor activity-defined regulons and transcription factor expression distinguishing in vitro and in vivo GPC4-subgroup hGPCs (Positive Log_2_FC indicates enrichment in in vivo hGPCs). **D-E.** Transcription factor regulon signaling networks of top active regulons in vitro (**D**) or in vivo (**E**) and their top targets (Triangles indicate regulons and circles downstream gene targets). **F.** Significant curated IPA terms differing between cultured and in vivo GPC4-subgroup hGPCs.

We next asked which transcriptional activators might be driving these significantly different transcriptional profiles. For this, we utilized pySCENIC trained on the in vitro and in vivo GPC4 population to first uncover potentially active transcriptional regulators ^36^. We then conducted a Wilcoxon rank sum test of each transcriptional regulator whose activity differed between the in vitro and in vivo GPC4s; this identified 278 significantly active regulons (FDR < 0.01, **Supplementary Data 4**). Of these, 36 were also found to be DE, with all but ASCL1 having concordant gene expression enrichment and activity (**Fig. 4C**). From these, we constructed a regulon signaling network for both cultured and engrafted GPC4 cells so as to identify their most critical transcriptional regulators as a function of in vitro vs in vivo residence – i.e., of the effects of transplantation on this critical stage of hGPC differentiation (**Figs. 4D-E**). In vivo, the key oligodendrocytic transcription factors SOX10 and OLIG2 were differentially active and expressed by GPC4s, along with a cohort of immediate early factors including JUNB, JUN, JUND, FOS, FOSB, EGR1, and EGR2 (**Fig. 4E**). These in turn were predicted to work in tandem to activate key downstream glial genes, including PDGFRA, PTPRZ1, CNTN1, TNR, BCAN, OLIG1, CCND1, and CSPG5. In contrast, cultured GPC4s expressed fewer actively coordinating transcriptional regulons (**Fig. 4D**), and these were characterized by regulons linked to younger and less-committed neural states, including DLX5, DLX2, DLX1, SOX2, SOX11 and SOX9. Critically, the expression and activity of these regulators declined after engraftment, leading to the repression of neural gene expression - which included DCX, STMN2, NNAT, CAMK2N1, MAP1B, and TAC3, from which few if any adventitiously-produced neurons could be identified.

We then utilized Ingenuity Pathway Analysis (IPA) to functionally profile these in vivo GPC4s, relative to their cultured counterparts (**Fig. 4F**). Among terms that were more active in vivo were those related to glial differentiation and myelination. These appeared to reflect the upregulation of TCF7L2, SOX10, STAT3, CTNNB1, FGFR1, KLF6, OLIG2, and THRA signaling by GPC4s. Furthermore, terms associated with heightened environmental interactions were similarly activated in vivo; these included cell-cell contact and semaphorin, integrin, NRG1, and EGR2 signaling. In contrast, cultured GPC4s expressed more active MYC and MYCN signaling, suggesting a less mature and more mitotically active state, along with greater pro-neural SOX11 signaling. These data indicate that transplantation and exposure to the in vivo tissue environment drives human GPCs to express a transcriptional signature distinct from and more differentiated than that achieved by hGPCs in vitro.

### Chromatin accessibility in vitro predicts differentiation bias in vivo

We next utilized our scATAC-Seq data describing the chromatin accessibility of cultured hGPCs (**Supplementary Fig. 3E**) to predict those GPC4 genes whose epigenetic state in vitro might have the most influence on cell fate after transplant. To do so, we fist intersected those genes DE between pre- and post-transplant GPC4s, with genes for which there was a consensus of gene expression and gene activity in vitro. This yielded genes that were both more accessible and differentially transcribed in cultured GPC4s, and which continued to be upregulated in vivo (**Supplementary Data 4**). These transcripts included CNTN1, TNR, CCND1, BCAN, PCDH17, ADGRL3, GRIA4, CA10 and NOVA1, as well as their predicted regulons, which included SOX10, OLIG1, and SOX8 (**Supplementary Fig. 5A**). In contrast, the downregulation of GPC4 gene expression in vivo followed the chromatin compaction and transcriptional suppression of GPC4 genes that first occurred in vitro including RTN1, GAD2, NEDD4L and BCL11A, as well as their predicted regulons DLX1, DLX2, DLX5, and FOXG1 (**Supplementary Fig. 5A**). Interestingly, we noted a subset of GPC4 genes whose chromatin became differentially accessible in vitro, but which did not exhibit differential RNA expression until later transplantation and in vivo residence (**Supplementary Fig. 4**). This latter set of genes appeared primed, but not actively transcribed until the cells encountered the in vivo environment; these genes included APOD, PCDH15, PLLP and GPR17, and their predicted regulons PRRX1, FOSB, EGR2, and ATF4 (**Supplementary Fig. 5B**). In contrast, chromatin was differentially compacted in GPC4 cells at loci that included CLVS2, DLX6, SYNPR, whose predicted regulons TEAD2 and PBX1 were differentially suppressed (**Supplementary Fig. 5B**). Together, these data suggest that the epigenetic state of cultured GPC4s significantly influenced the potential scope of their downstream gene expression, thus biasing the differentiated fate of these cells after transplantation.

### Resident mouse brain cells drive the specification and differentiation of human GPCs in vivo

Given the substantial transcriptional differences between cultured hGPCs and their transplanted, tissue-resident counterparts, we next sought to identify potential environmental drivers of this process. To do so, we first classified the murine cells that were present in our captures of hGPC-engrafted callosal white matter (**Fig. 5A**). Following integration and leiden clustering, we identified 10 major cell types including mouse GPCs, imOLs and maOLs, astrocytes, microglia, endothelial, pericytes, NPCs, macrophages, and ependymal cells (**Figs. 5A-B, Supplementary** Figs. 6A-B**, Supplementary Data 5**). Notably, mouse GPCs were nearly absent from these captures, likely due to their replacement by engrafted, competitively dominant hGPCs^37^.

**Figure 5.**
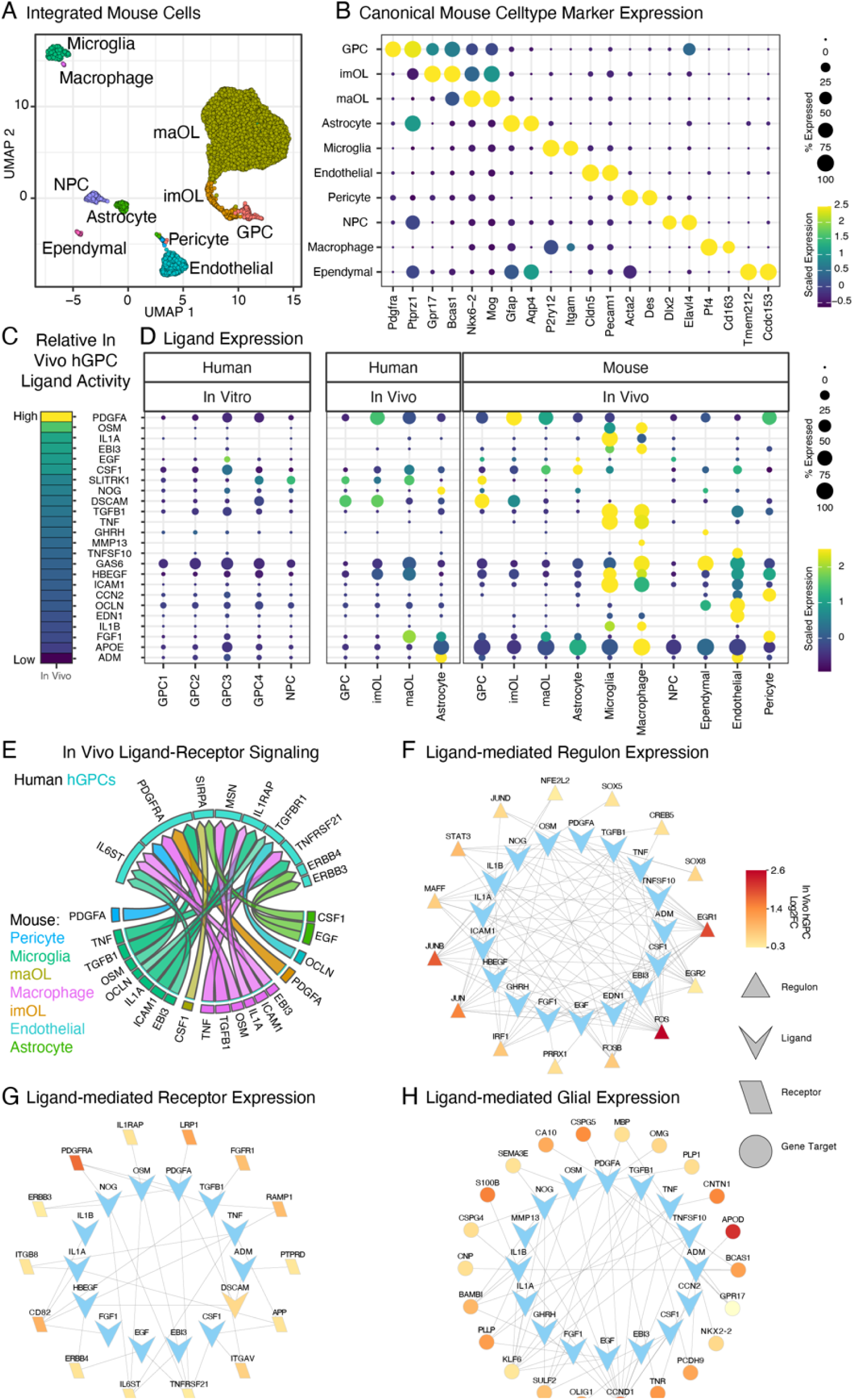
Cell-cell interactions drive hGPC differentiation in vivo. **A.** Integrated UMAP plot of mouse cells from hGPC engrafted shiverer white matter. **B.** Dot plot of canonical cell type markers. **C.** Relative NicheNet ligand activity of in vivo vs in vitro hGPCs and **D.** their scaled gene expression in all cultured human populations, and in all major human and mouse brain populations in vivo. Ligands were selected based on high in vivo vs in vitro hGPC activity, in vivo ligand expression, and in vivo hGPC receptor expression. **E.** Chord plot of top mouse ligand-human receptor pairs differentially active in hGPCs in vivo. **F-H.** Top ligand-mediated expression networks active in hGPCs in vivo, for regulon transcription factors (**F**), receptors of curated ligands (**G**), and glial genes (**H**).

We next used data derived from cultured hGPCs prior to transplant, as well as from the in vivo mouse and human cells, to predict hGPC ligand-receptor interactions differentially active as a function of context, using the NicheNet package ^38^. We focused on hGPCs classified **in Fig. 2D** as GPC4 as receiver cells, since this population comprised the most hGPC-specified subpopulation, both in vitro and in vivo (**Fig. 4A**). Displayed ligands were curated based on a composite of higher ligand activity (**Fig. 5C, Supplementary Data 5**), higher cell type-selective expression of the ligand (**Fig. 5D, Supplementary Data 5**), and sufficient receptor expression by hGPCs in vivo (>10% of cells) as well as, in many cases, higher receptor expression (**Supplementary Fig. 6C, Supplementary Data 5**).

From this analysis, the canonical hGPC specifying ligand PDGFA was predicted to exert the strongest transcriptional downstream influence on hGPCs in vivo (**Fig. 5C**) ^39,40^. Human and mouse oligodendrocytes – both immature and mature - were the highest PDGFA ligand-expressing sender cells in vivo, signaling primarily through PDGFRA, the most significantly upregulated receptor among hGPCs in vivo, when compared to their in vitro counterparts (**Figs. 5D-E**, **Supplementary Fig. 6C**). Microglia and perivascular macrophages were also predicted to express numerous active ligands, including OSM, IL1A, EBI3, TGFB1, TNF, MMP13, GAS6, HBEGF, ICAM1, IL1B, and APOE^41–43^. These in turn were predicted to signal through IL6ST (Also known as GP130), IL1RAP, TGFBR1, TNFRSF21 and TNFRSF1A, LRP1 and LRP6, as well as via the integrins ITGB8, ITGB1, and ITGAV, among others. Mouse astrocytes were the primary source of the ligands EGF and CSF1, signaling through ERBB3/B4 and SIRPA respectively, whereas human astrocytes expressed relatively high levels of NOG and ADM, which influenced hGPC fate via BMP pathway suppression or RAMP1 and RAMP2 activity, respectively. In parallel with these pathways, mouse endothelial cells were predicted to direct hGPC activity in vivo via TNFSF10/TRAIL, EDN1, and ADM.

On the basis of these data, we constructed ligand-target networks from those ligands predicted to be more active in hGPCs in vivo relative to their counterparts in vitro (**Figs. 5C-D, Supplementary Data 5)**. Our targets included predicted transcriptional regulons active in hGPCs in vivo (**Fig. 5F**, from **Fig. 4C**), receptors of curated ligands (**Fig. 5G**, from **Supplementary Fig. 6C**), and salient developmentally-regulated glial genes (**Fig. 5H**, from **Fig. 4B**). By this means, we predicted a set of transcriptional regulons differentially active in transplanted hGPCs that included SOX10, SOX8 and OLIG2, as well as EGR1, EGR2, STAT3, and the intermediate-early activators FOS, FOSB, JUN, JUNB, JUND and MAFF (**Fig. 5F**). Driving these, we predicted that PDGFRA, ERBB3, ERBB4, IL6ST, LRP1, and FGFR1 comprised those DE hGPC receptors most regulated via ambient expressed ligands, whether of human or mouse origin (**Fig. 5G**). Furthermore, a number of maintenance-associated hGPC genes were predicted targets of in vivo-enriched ligands, as were genes linked to oligodendrocytic fate restriction; these respectively included NKX2-2, OLIG1, SULF2, CSPG4, CSPG5, GPR17, BAMBI, TNR and CCND1 regulating hGPC maintenance; and BCAS1, PLP1, CNP, MBP, and CNTN1 directing initial oligodendrocytic differentiation. Thus, transplanted hGPCs are exposed to a variety of ligands in vivo to which they are not exposed in vitro, whose integrated signal output serves to drive their context-dependent maturation in vivo.

### Downstream regulators drive human oligodendrocytic maturation and myelination in vivo

We next utilized these data to identify transcriptional regulators driving further in vivo human oligodendrocytic maturation and myelination in vivo, during their sequential maturation through imOL and maOL stages. To do so, we first identified differentially expressed genes between each in vivo transition (MAST, FDR < 0.01, log_2_FC > 0.25, imOL vs GPC 1,086 genes, maOL vs imOL 1,549 genes, **Supplementary Data 6**). We then ran pySCENIC on the human cells and similarly tested for differential regulon activity between the same cell type transitions (Wilcoxon rank sum test, FDR < 0.01, imOL vs hGPC 309 regulons, maOL vs imOL 277 regulons, **Supplementary Data 6**). Notably, the gene expression and regulon activity of top transcription factors were tightly coordinated across glial differentiation when plotted in pseudotime (**Fig. 6A**, pseudotime from **Fig. 3A**). We then constructed regulon signaling networks for each oligodendroglial stage transition, which served to emphasize the most distinct stage-selective transcriptional features of each cell state (**Figs. 6C-D**, top differentially expressed targets plotted across pseudotime in **Fig. 6B**).

**Figure 6.**
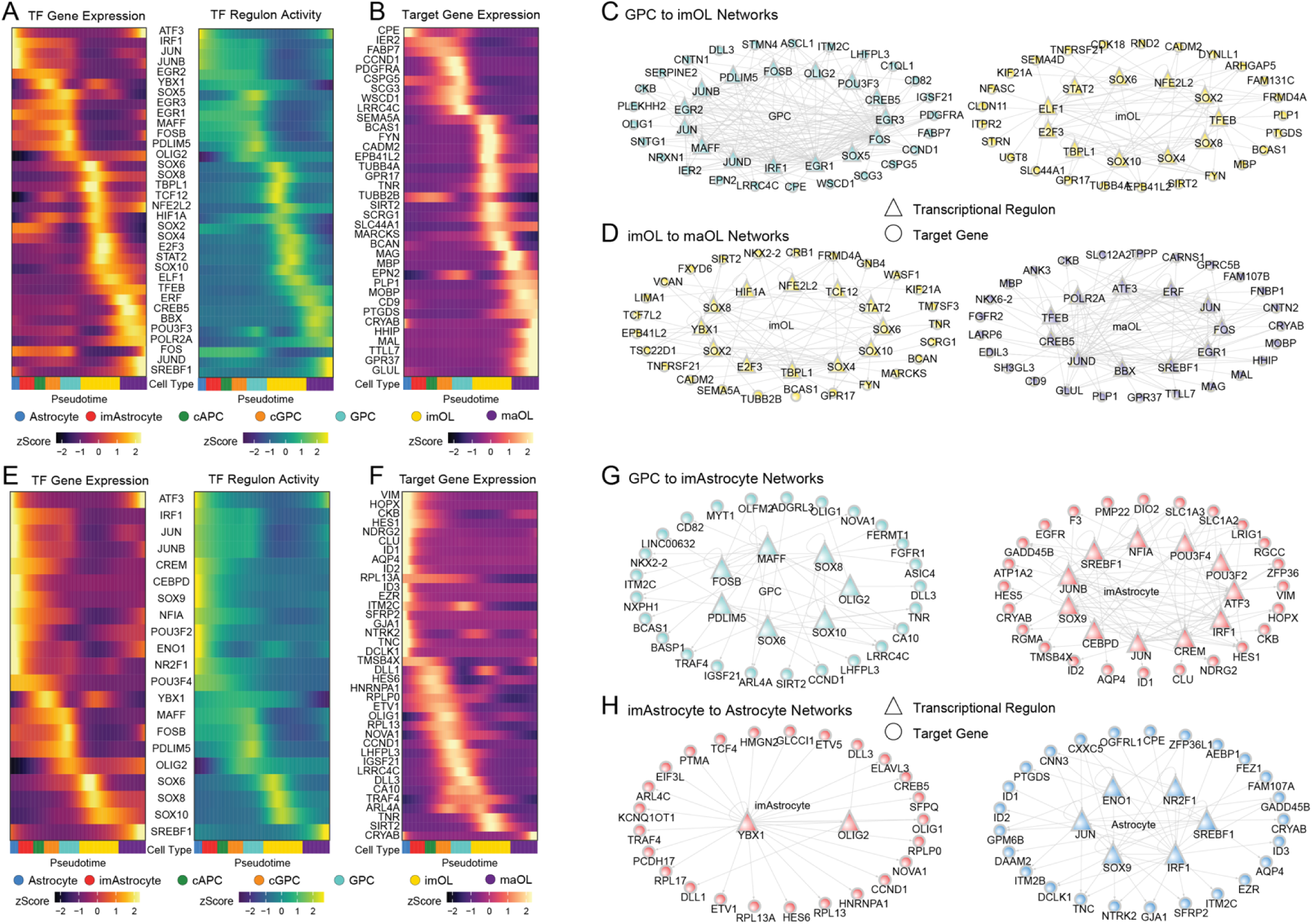
Key transcriptional regulators drive human oligodendrocyte and astrocyte differentiation in vivo. pySCENIC activity and gene expression of top oligodendroglial (**A**) and astrocytic (**E**) lineage transcription factor (TF) regulons plotted in pseudotime (slingshot^34^). Regulons were determined via enrichment, of adjacent glial lineage cell type regulons of the most differentially expressed regulon targets. Top oligodendroglial (**B**) or astrocytic (**F**) lineage regulon target genes. **C-D, G-H.** Top regulon signaling networks enriched in **C**) hGPCs relative to immature oligodendrocytes (imOL), **D**) imOLs vs mature oligodendrocytes (maOL), **G**) GPCs vs immature Astrocytes (imAstrocytes), and **H**) imAstrocytes vs mature astrocytes.

Within the hGPC vs immature oligodendrocyte gene regulatory network, numerous transcriptional regulators were observed to define the progenitor state; these included OLIG2, SOX5 and POU3F3/BRN1, as well as EGR1, EGR2, EGR3, and MAFF (**Fig. 6C** left). Interestingly, many of these were among those whose activity rises in vivo, compared to cultured hGPCs (**Fig. 4C**). Among hGPC enriched genes that were activated by these regulons were numerous early glial genes, including PDGFRA, NFIA, CCND1, CSPG5, NRXN1, OLIG1, CNTN1, and ASCL1. PDGFRA was the most GPC enriched gene, and was predicted to be directly regulated by EGR1, EGR3 and SOX5. Yet at the onset of differentiation, immature OLs upregulated the activity of SOX10 as well as SOX8, SOX6, SOX4, and TFEB, among others (**Fig. 6C** right). These regulons were then predicted to drive the early expression of myelin and oligodendrocytic genes, including MBP, BCAS1, PLP1, CLDN11, UGT8, GPR17 and SIRT2, and SEMA4D. Of these, TFEB was predicted to directly target the top 4 most upregulated genes during the beginning of oligodendrocyte differentiation: PLP1, PTGDS, BCAS1, and MBP.

As oligodendrocytes matured, they downregulated their expression of transcriptional regulons including SOX10, SOX8, SOX6, SOX4 and NFE2L2 (Also known as NRF2), all of which were associated with the initiation of oligodendroglial differentiation (**Fig. 6D** left). This in turn led to the downstream reduction of early oligodendroglial maturation-associated genes, that included BCAS1, GPR17, FYN, BCAN, TNR, WASF1, NKX2-2, SIRT2, and TCF7L2. In contrast, TFEB activity continued to increase in mature oligodendrocytes, while ATF3, ERF, SREBF1, and BBX activities all rose along this trajectory (**Fig. 6D** right), as did NKX6-2, MAL, GPR37 and CARNS1, as well as the myelin genes MAG, MOBP, PLP1 and MBP. Together, these data highlight the complex orchestration of transcription factor-driven signaling cascades by which oligodendrocytic differentiation from PSC-derived hGPCs is achieved in vivo.

### A distinct set of transcriptional regulators direct human astrocytic differentiation in vivo

We next explored astrocytic differentiation from hGPCs in our chimeric white matter using the same approach as employed for the oligodendrocytic lineage. This analysis yielded 881 genes DE between immature astrocytes and GPCs, and 630 between immature and mature astrocytes (MAST, FDR < 0.01, log_2_FC > 0.25, **Supplementary Data 7**). These gene expression changes were predicted to be orchestrated by 244 regulons during the transition from GPCs to imAstrocytes, and 155 during the progression to mature astrocytes (Wilcoxon sum rank test, FDR <0.01, **Supplementary Data 7**). Just as with the oligodendrocytic lineage, transcription factor expression and predicted activity were concordant in pseudotime across astrocytic differentiation (**Fig. 6E**). We next constructed regulon signaling networks regulating the transition of GPCs to immature astrocytes and then to mature astrocytes (**Figs. 6G-H**, top DE targets plotted across pseudotime in **Fig. 6F**).

The same top transcriptional regulators identified in the GPC to immature astrocyte transition were also observed during the early stages of the oligodendrocyte trajectory; these included OLIG2, SOX10, SOX8, SOX6, MAFF, FOXB, and PDLIM5 (**Fig. 6G** left), suggesting that they remained poised for either astrocytic or oligodendrocytic differentiation at this stage. Yet as these progenitors began to differentiate as immature astrocytes, they increased their expression and activity of SOX9, SREBF1, NFIA, and CEBPD (**Fig. 6G** right), with further downstream activation of SLC1A2, SLC1A3, AQP4, ID1, ID2, CLU, HES1, HES5, HOPX, VIM, F3, and EGFR. At the same time, OLIG2 and YBX1 signals sharply declined with terminal astrocytic maturation (**Fig. 6H**, left), leading to the decreased expression of their downstream oligodendrocytic signatures. At the same time, SOX9, SREBF1, IRF1, and JUN signaling all increased, pare passu with the induction of the ENO1 and NR2F1 regulons (**Fig. 6H** right). This process drove the upregulation of a cohort of mature astrocytic genes, including AQP4, GJA1 (Also known as CX43), TNC, EZR and CRYAB. Of note, a number of transcripts were associated with terminal differentiation of hGPCs towards *either* astrocytic or oligodendrocytic fate. For instance, SREBF1 expression was predicted to enable differentiation along both astrocytic and oligodendrocytic paths, highlighting the importance of cholesterol-regulated pathways in glial differentiation ^44^. Together, these networks serve to describe the tissue-driven signaling networks triggered in transplanted hGPCs that direct their differentiation beyond that achieved in vitro, and by which cell fate and state may be regulated and stabilized after transplantation.

## DISCUSSION

Human GPCs comprise a potential cell therapeutic for the treatment of a variety of neurological diseases caused or exacerbated by glial pathology ^7,8,45,46^. We sought here to better define both the composition and epigenetic regulators driving hGPC differentiation in vitro, and to explore how transplantation of these progenitors into the murine brain microenvironment shape their further differentiation into mature oligodendrocytes and astrocytes^47–51^. To this end, we used a combination of scRNA-Seq, ATAC-Seq and CUT&TAG assessment of stage-specific histone modifications to define the cell composition at various points of glial differentiation by both ESC- and iPSC-derived hGPCs, generated using our in vitro differentiation protocol ^11^. We found that our GPC differentiation protocol produced cells free of pluripotent signatures, thus mitigating a known risk of cell therapy^52^. While pluripotent cells highly expressed the canonical PSC markers LIN28A, POU5F1 and APELA, these genes were undetectable by the earliest appearance of the GPC stages ^14,53^. Furthermore, no GPCs assayed in vitro, nor any human cells extracted back from chimeric mice co-expressed these factors, indicating that differentiation eliminated PSCs from our preparations, at least to our limits of detection, providing reassurance regarding their safety as potential clinical therapeutics.

We further characterized this in vitro differentiation process epigenetically using CUT&Tag-seq, focusing on three histone marks linked to glial development and function: H3K4me3, H3K27ac, H3K27me3 ^54–60^. We observed cooperation of these marks in promoting pluripotent gene expression at the PSC stage, but remodeling of the chromatin landscape during in vitro differentiation, leading to the repression of the pluripotent signature and the activation of a glial program. This process may have been in part orchestrated by EZH2, known to trimethylate H3K27 which was found to be more highly expressed at the in vitro hGPC stage, consistent with previous findings highlighting its importance in glial specification^61^. CHD7 was also upregulated during in vitro hGPC differentiation; it has been implicated in promoting myelinogenic gene expression through cooperation at H3K27ac bound enhancers^62^, and via modulating chromatin accessibility at genes such as GPR17, SOX10, and SIRT2, each of which we found to be differentially accessible in our hGPC scATAC-Seq data^63^.

scRNA-seq revealed the composition of pre-transplant cultured hGPCs to be that of NPCs, four populations of early GPCs, and a small percentage of immature oligodendrocytes. Among these populations were GPC2, characterized by cycling activity, GPC3, enriched for astrocytic expression, and GPC4, which exhibited the highest expression of genes associated with oligodendroglial progenitors, including PDGFRA, OLIG2, OLIG1, SOX10, and NKX2-2. Transcriptional hallmarks of these populations as detailed above are largely shared with tissue-derived fetal human ^64,65^ and murine progenitors^35,66,67^ alike. Integrated analysis with human glia extracted back from the white matter of 19-week-old shiverer chimeras indicated that in vitro GPC4 progenitors clustered with in vivo GPCs, suggesting their relative similarity. Furthermore, fate trajectory analysis similarly predicted that in vivo GPCs emerged from cultured GPC4s via GPC1-3. Interestingly, this trajectory analysis predicted both oligodendrocytic and astrocytic differentiation from GPC4s, suggesting the in vivo derivation of astrocytes from this common progenitor, rather than from the more astrocyte-biased in vitro pool of GPC3s. Supporting this prediction, cycling progenitors in vivo consisted of two populations, that were respectively enriched in early oligodendroglial and astroglial genes, each of which was predicted to mature along its eponymous lineage. That said, further lineage analyses will be necessary to assess the phenotypic plasticity and malleability thereof of these progenitor cell states in vivo.

Importantly, despite the transcriptional similarities and co-clustering of cultured GPC4 cells and GPCs in vivo, these two populations nonetheless exhibited clear differences in their expression profiles. In vitro, GPC4 GPCs were enriched for genes associated with fetal human GPCs, including BCL11A, TEAD2, SOX4, SOX9, SOX2, ETV1, HDAC2, and TET1, along with early neural identity markers^35,68^. In contrast, once in vivo, the hGPCs downregulated this neural signature in favor of canonical oligodendroglial genes, including PDGFRA, OLIG2, OLIG1, SOX10, and PTPRZ1^24,31,69,70^. Our analysis predicted that this process was driven via the upregulation and increased activity of the oligodendrocytic transcription factors SOX10 and OLIG2, downstream of a cohort of immediate early factors, and accompanied by the suppression of early neuronal transcripts, including DLX1, DLX2, DLX5, and SOX11.

These differences in in vitro vs. in vivo gene expression by GPC4-cluster GPCs appeared driven by the host white matter environment, in which the murine astrocytes, pericytes, endothelial cells, microglia and perivascular macrophages all expressed potentially-consequential ligands. This contrasted with the relatively weak downstream ligand activation of cultured GPC4s by other glia in vitro. Of note though, our data predicted that PDGFA secretion from both murine and human oligodendrocytes, would elicit an upregulation of PDGFRA receptor expression by GPCs, the activation of which would then drive the expression and activity of EGR1, EGR2, and SOX8 regulons ^21,71,72^. Indeed, EGR1 and SOX8 have been shown to be important in early GPC specification ^73,74^, and were predicted by our data to be upstream of the critical glial genes SOX10, OLIG1, OLIG2, BCAN, PLP1, BCAS, and CNTN1, in addition to sustaining PDGFRA expression.

Astrocytes interact with hGPCs to potentiate oligodendrocytic maturation and myelination, so that astrocytic dysfunction can lead to oligodendroglial loss, such as that seen in Alexander disease and vanishing white matter disease^46,75–79^. Transplantation of cultured GPCs gives rise to astrocytes as well as oligodendrocytes which - in this chimeric context and timeframe - co-exist with murine astrocytes^12,48,50^. We thus investigated which astrocytic ligands might influence in vivo GPC maturation. We found that astrocytes – whether of mouse or human origin - were predicted to strongly influence engrafted hGPCs via ligands encoded by ADM, NOG, CSF1, and EGF^80–82^. Among these, NOG was of particular interest, since the BMPR2 and BMPR1A receptors – whose activities are inhibited by NOG-dependent inhibition of their BMP ligands - were upregulated on GPCs in vivo, and their signaling was predicted to increase SOX5 and PRRX1 transcriptional regulon activity. The latter genes have been linked to glial maturation ^18,83,84^, suggesting the astrocytic, NOG-dependent moderation of hGPC differentiation. Similarly, the EGF pathway receptors ERBB3 and ERBB4 were both upregulated in GPCs in vivo^64,82,85^, again suggesting local astrocytic regulation of oligodendrocytic maturation by GPCs.

This analysis also predicted the potential importance of a number of host microglial and perivascular ligands in oligodendrocytic production and/or maturation. Our analysis identified microglial ligands of GPC receptors that include OSM, GAS6, TGFB1, TNF, and HBEGF ^41–43,86–89^. Downstream of OSM, the transcriptional regulons EGR1 and EGR2 were upregulated, likely reflecting the activation of IL6ST, the oncostatin M (OSM) receptor, in hGPCs in vivo. Similarly, microglial TGFB1 and TNF were both predicted to upregulate the GPC expression of the EGR1, EGR2 and STAT3 regulons, the transduction of which was likely amplified by the GPC expression of the TNF receptors TNFRSF1A and TNFRSF21. In addition, microglial HBEGF was predicted to signal via the hGPC-enriched receptors CD82, ERBB4, and CD9, whose activation has been linked to the appearance of oligodendroglial genes including PDGFRA, GPR17, MBP, PLLP and CNTN1 ^24,90,91^. Yet microglia are not alone among non-neural regulators of GPC fate; vascular endothelial and pericyte ligands have also been linked to the behavior of GPCs ^92–94^. We found that ADM, encoding adrenomedullin, was expressed by endothelial cells – just as it is by astrocytes - thus reinforcing the possibility of an ADM to RAMP2/RAMP3 signaling axis regulating the interaction of GPCs with the local environment. Similarly, endothelial EDN1, previously reported to mediate experience-dependent myelination, was also predicted to signal through the GPC-enriched EDNRB^95^. Together, these data highlight the potential import of both microglial and vascular interactions with GPCs in the regulation of progenitor maturation and myelination ^64,88,96^.

Together, these data provide a map of the transcriptional networks and epigenetic controls associated with hGPC diversification and differentiation, both in vitro and in vivo. Our data highlight the role of the tissue environment on that process, by identifying the contextual cues and signaling networks that underpin the differentiation of hGPCs after their introduction into host brain. As such, these data provide us insight into the responses of hGPCs to transplantation, and hence into the behavior of these cells in the potential clinical scenario of their transplantation into subjects with glial dysfunction or myelin loss, as in the childhood leukodystrophies or adult autoimmune dysmyelinations. By demonstrating the elimination of persistent undifferentiated cells during the differentiation of GPCs from hESCs, and the further context-dependent terminal differentiation of these cells after their transplantation into the tissue environment, our findings suggest the safety of hGPCs as potential cellular therapeutics ^8,97^, while also identifying genes and pathways whose modulation may enable gene therapeutic strategies towards the treatment of glial disease.

## METHODS

### Human Cell Lines

The embryonic stem cell line WA09 was purchased from WiCell. The induced pluripotent stem cell line C27 was generously provided to us by Lorenz Studer ^11,117^.

### Mouse Models

Homozygous MBP^shi/shi^ shiverer mice (The Jackson Laboratory) were bred with homozygous Rag2^-/-^ immunodeficient mice on the C3h background (Taconic) to produce myelin-deficient and immunodeficient mice ^118^. Mice were housed in a temperature and humidity-controlled environment (64-73°F, 30-70% humidity) within a pathogen-free colony room on a 12:12-hour light cycle. They were given ad libitum access to Modified ProLab RMH 3000, 5P00 containing 0.025% trimethoprim/0.124% sulfamethoxazole (Mod LabDiet 5P00), and autoclaved acid water (pH 2.5-3.0).

### Production of GPCs from human iPSCs or ESCs

Human induced pluripotent stem cells (C27 ^117^) or embryonic stem cells (WA09) were differentiated into GPCs using our previously described culture conditions and preparation protocol ^11,48,50^. Briefly, cells were serially differentiated to neuroepithelial cells, then pre-GPCs and GPCs. The latter were typically apparent by 70 days in vitro, and expanded with progressive in vitro enrichment until harvest for transplantation, between 130-160 DIV.

### FACS of in vitro cells for scRNA-seq capture

Unsorted PSC and GPC stage cells_were incubated in Accutase (Millipore Sigma Cat. A6964) diluted 50% in DPBS for 6 minutes at 37 degrees. Cells were resuspended in DMEM and filtered for FACS. Cells were selected based on forward and side scatter, in addition to negative gating of 4′,6-diamidino-2-phenylindole (DAPI, 1LµgLml^−1^; Thermo Fisher Scientific, D1306) staining to remove dead cells and debris. Cells were then captured for scRNA-seq on a 10X Genomics chromium controller.

### Generation of human glial chimeric mice

Homozygous shiverer (MBP^shi/shi^) x Rag2^-/-^ mice were perinatally transplanted (at P0 or P1) with 3 x 10^5^ hESC or iPSC-derived hGPCs, delivered into anterior and posterior callosal anlagen bilaterally, as previously described ^13,23,48,50^.

### Immunohistochemistry

Human glial chimeric mice were euthanized with euthasol at 19 weeks and perfused intracardially with HBSS containing magnesium chloride and calcium chloride followed by 4% paraformaldehyde in 0.1 M PBS. Brains were extracted, post fixed in 4% paraformaldehyde for 2h, and stored in 1X PBS. Brains were cryopreserved in 30% sucrose diluted in 0.1M PBS before OCT embedding. Brains were sectioned sagitally at a thickness of 20 μm on a Leica Cryostat. Sections were blocked in 5% normal goat serum in 0.1 M PBS and 0.3% Triton-X for 1h at RT and incubated for two days at 4°C for huNA (1:800, mouse, Millipore), huGFAP (1:500, mouse, BioLegend), or MBP (1:25, rat, Abcam). AlexaFluor (Thermo Fisher) conjugated secondary antibodies were diluted 1:400 in blocking buffer for 1h at RT. Sections were stained with DAPI for 5 min, coverslipped, and imaged on a Leica DM6000B confocal microscope.

### FACS isolation of chimeric white matter cells for scRNA-Seq

Human and mouse cells were isolated from 19 week shiverer mice as previously described^12,23^. Briefly, chimeras were euthanized with euthasol and perfused intracardially with HBSS containing magnesium chloride and calcium chloride. Their brains were removed and immersed in ice-cold HBSS and their corpora callosa dissected and chopped into small pieces. This tissue was then incubated in a papain/DNase dissociation solution at 37 °C for 50 min. Papain was inactivated with the addition of ovomucoid dissolved in EBSS. A single-cell suspension was then achieved via repeated pipetting. Cells were pelleted, resuspended into MEM, and filtered for FACS. Cells were selected based on forward and side scatter in addition to negative gating of 4′,6-diamidino-2-phenylindole (DAPI, Thermo Fisher Scientific, D1306) staining to remove dead cells and debri (1LµgLml^−1^). Cells were then captured for scRNA-seq on a 10X Genomics chromium controller.

### Alignment and preprocessing of scRNA-seq samples

Samples produced using 10X Genomics scRNA 3.1 Chemistry were aligned STARsolo ^119^ with the following flags: “--twopassMode Basic --outSAMtype BAM Unsorted --readFilesCommand zcat --soloType CB_UMI_Simple --soloCBwhitelist 3M-february-2018.txt --soloUMIlen 12 --soloUMIfiltering MultiGeneUMI -- soloCBmatchWLtype 1MM_multi_pseudocounts --limitSjdbInsertNsj 2000000”. Samples produced using 10X Genomics scRNA 2.0 chemistry were aligned using the following STARsolo flags: “--twopassMode Basic --outSAMtype BAM Unsorted --readFilesCommand zcat --soloType CB_UMI_Simple -- soloCBwhitelist 737K-august-2016.txt --soloCBstart 1 --soloCBlen 16 --soloUMIstart 17 --soloUMIlen 10 -- soloBarcodeReadLength 0 --soloUMIfiltering MultiGeneUMI --soloCBmatchWLtype 1MM_multi_pseudocounts --limitSjdbInsertNsj 2000000.” All samples were aligned to a custom dual species reference using STARsolo. This reference was constructed by concatenating GRCh38 Ensembl 106 to GRCm39 Ensembl 106 and filtering for transcript types: “protein_coding”, “lncRNA”, or “miRNA”. Following alignment, the STARsolo filtered UMI count matrix was split into components containing either mouse or human genes in R^98^. For each species, the respective count matrix was filtered for cell barcodes that possessed at least 500 unique genes expressed and less than 15% mitochondrial gene expression in Seurat for further analysis^99^.

### Integration of multiple samples and dimensionality reduction

Raw UMIs from cells passing the cut-off criteria in all samples were merged into a matrix, and used as input for data integration. Data were integrated and accounted covariates including cell line, 10X chemistry, percent mitochondrial gene expression, depth, and complexity with scVI^114^. Cellular architecture and heterogeneity were visualized via UMAP ^120^, and leiden cell clusters determined from integrated model normalized values using scanPY ^113^ with default parameters. Identity of each cluster was manually assigned based on the canonical cell type markers shown in the corresponding figures and Supplemental Tables.

### Differential expression analysis

For differential expression, gene sets were first filtered for appreciably expressed genes (10% detection in any comparison group). Differentially expressed genes were identified using MAST with cell line, chemistry version, and scaled number of unique features as fixed effects and capture as a random effect^16,121^. Multiple comparisons were controlled using the false discovery method. For differential expression between in vivo and in vitro GPCs, genes that were identified as having infinite-fold changes between groups from the MAST package were set to the max absolute log_2_ fold change ± 0.1 for visualization and downstream inclusion.

### Bulk CUT & Tag of human PSCs or GPCs

Pluripotent stage cells (0 DIV) were harvested by incubation in Gentle Cell Dissociation Reagent (STEMCELL Technologies Cat. 07174) for 7 minutes at room temperature. GPC stage (180 DIV) cells were incubated in Accutase (Millipore Sigma Cat. A6964) diluted 50% in DPBS for 6 minutes at 37 degrees. After centrifugation, cells were resuspended in gentle nuclei isolation buffer (10mM HEPES pH 7.9, 10mM KCl, 0.5mM Spermidine, 0.10% IGEPAL, 1X cOmplete EDTA-free protease inhibitor cocktail (Roche Cat. COEDTAF-RO)).

Following nuclei extraction, buffers were prepared and Cleavage Under Targets & Tagmentation (CUT & Tag) was conducted following the CUTANA^TM^ CUT & Tag protocol (EpiCypher, version 1.7 ^20^). Samples included: PSCs: n=4 for each histone mark for WA09, and n=2 H3K4me3 and H3K27ac, n=1 for H3K27me3 for C27; GPCs: n=2 for WA09, and n=3 for C27 for each histone mark. In brief, nuclei were adsorbed to Concanavalin A Conjugated Paramagnetic beads (EpiCypher Cat. 21-1401) and incubated with rabbit anti-H3K4me3 (Cell Signaling Technology Cat. 9751S), H3K27me3 (Cell Signaling Technology Cat. 9733S), or H3K27ac (Cell Signaling Technology Cat. 8173S) overnight at 4°C. The following day, anti-rabbit secondary antibody (EpiCypher Cat. 13-0047) was added, followed by pAG-Tn5 (EpiCypher Cat. 15-1117) to induce tagmentation of the chromatin. Tagmented DNA was indexed and amplified for sequencing, then sequenced on the NextSeq 550 System using the Mid-Output Kit (Illumina).

### Bulk CUT&Tag Analysis

Sequencing data were aligned to GRCh38 using bowtie2 (version 2.3.5.1) with the following settings: [--local --very-sensitive --no-mixed --no-discordant --phred33 -I 10 -X 700]^105^. Reads were filtered with samtools (version 1.9) with the following settings [-F 4 -q 30], and the duplicates were removed with picard (version 3.0.0)^106,107^. After filtration, bedgraphs were generated using bedtools (version 2.30.0), which were then used to call peaks with SEACR (version 1.3), using a threshold of 0.01 and stringent settings^108,109^.

The peak files and associated bam files were used as input for DiffBind (version 3.12.0,) with default settings, to identify differential peaks between GPC and Pluripotent stage using cell set as a covariate (FDR<0.05)^101^. The differential peaks for H3K4me3 and H3K27me3 were annotated to gene promoters using ChIPseeker (version 1.38.0) with the promoter region set to be within a 2kb+/- window of the transcription start site (TSS)^102^. The differential peaks for H3K27ac were annotated to putative enhancers using the GeneHancer database using bedtools^122^. To identify concordant patterns of transcriptional and epigenetic profiles, differentially expressed genes were identified via scRNA-seq of v3.1 libraries using MAST of pluripotent vs GPC cells with line and number of unique genes as fixed covariates and capture as a random effect (FDR adjusted p-value < 0.01, log2FC > 0.25)^16^. Genes were included if they were expressed at either stage in >30% of cells and if they had an average expression at either stage of >0.3 UMIs/10,000.

To plot the H3K4me3/H3K27me3 profile at promoters and H3K27ac profile at putative enhancers for significantly up/down-regulated genes, normalized bigwig files were generated using deeptools bamCoverage (version 3.5.1) with the following options [--binSize 10 --normalizeUsing RPKM -- extendReads], and then used as input for deeptools computeMatrix and plotHeatmap to generate the plot^110^. The gene annotations were obtained from Ensembl Release 106, and the putative enhancers were identified using the GeneHancer database with bedtools. The genome browser snapshots were prepared with Integrative Genomics Viewer (IGV) (version 2.8.13)^111^.

### scATAC-seq of human GPCs

hESC-derived GPCs (WA09) were dissociated at 180 DIV with 50% Accutase in Dulbecco’s PBS, and nuclei isolated according to the 10X Genomics Chromium 3’ v1.1 scATAC kit. After library production, samples were sequenced on an Illumina NovaSeq6000 at the University of Rochester’s Genomics Resource Center.

Sequencing data were aligned to GRCh38 using Cell Ranger ATAC (version 2.0.0) with default settings^32^. Counts, metadata, and fragments were imported into R using Seurat (version 4.3.0) and Signac (1.14.0)^99,115^. Peaks were filtered to standard chromosomes and annotated to Ensembl 106 using a modified version of GetGRangesFromEns that returns all biotypes. Nuclei were scored for TSS enrichment, fraction of reads in peaks, and blacklist ratio. Nuclei were then filtered for quality leaving 17,649 nuclei (Peaks > 9000 and < 100,000, percent reads in peaks > 40, blacklist ratio < 0.01, nucleosome signal < 4, and TSS enrichment > 4). The term-frequency inverse-document-frequency was computed, top features calculated (min cutoff = q0), and singular value decomposition run. Gene activities were calculated using Signac’s GeneActivity function on all genes and then log normalized. Predicted labels from the in vitro scRNA subpopulations were transferred using FindTransferAnchors with CCA reduction on all genes that were significantly enriched in any cultured GPC subpopulation from the scRNA-seq (**Supplementary Data 2**) and then TransferData using the LSI reduction on dims 2:30. A UMAP plot was generated within Seurat (LSI reduction, min.dist = .9, dims = 2:30). Significantly enriched gene activities in each cultured GPC subpopulation were determined using the wilcoxauc function (Adjusted p-value < 0.01) from the Presto package (version 1.0.0)^116^. Immature oligodendrocytes were not tested for accessibility or used for plotting due to their rarity (6 nuclei). Log2 fold changes of top enriched genes in each cultured GPC subpopulation were plotted with ggplot2 with their differential enrichment gene expression shown below^100^. A scatter plot was constructed using the log2 fold changes of cultured GPC4 differential gene expression enrichment and log2 fold changes of cultured GPC4 differential gene activity enrichment with assay significance indicated by color. Differential accessibility of loci was also calculated using the wilcoxauc function on scATAC-seq peaks (Adjusted p-value < 0.01). Curated loci were plotted using Signac’s CoveragePlot function (+/− 1,000bp).

### Identification of transcription factor regulons

Genes were first filtered to retain only those that expressed at least 3 counts in at least 1% of the cells. The filtered raw matrix was then used as input for the standard pipeline of pySCENIC^36^ to identify each transcription factor and its putative downstream targets in our data set. These gene sets are referred to as regulons and are assigned “Area Under the Curve” (AUC) values to represent their activities in each cell, with higher values indicating higher predicted activity. Wilcoxon rank sum tests were used to determine differential regulon activity in Seurat (FDR < 0.01). pySCENIC is not effective at identifying repressive relationships^36^ so transcription factors whose primary regulatory mechanisms are repressive in the literature were removed prior to mining downstream targets. Networks were constructed of regulons and their targets based on concordant directionality of regulon activity and that transcription factor’s gene expression in addition to concordant target gene expression directionality. The top 25 most differentially expressed target genes by log2 fold change, that were targeted by a transcription factor that targeted at least 2 genes of these 25, were selected for display excluding other already included regulon genes.

### Pseudotime analysis

Pseudotime analysis was conducted either with palantir^33^ or slingshot ^34^. For Palantir, depth normalized values from the scVI integrated model of GPC stage in vitro and chimeric in vivo cells were obtained for the top 3,000 highly variable genes. The standard Palantir pipeline was then followed using 10 components for the diffusion map step and 5 eigen vectors for determining the multiscale space. The starting cell was selected based on the highest DLX5 expression and the terminal cells were selected based on the highest expression of MOBP, AGT, and NSG2 for oligodendrocytes, astrocytes, and NPCs respectively. Trajectories were obtained in python via the plot_trajectory function and superimposed on UMAP plots generated in R^98,112^. Branch probability plots were generated with ggplot2 from values obtained in Palantir.

For slingshot pseudotime analysis of in vivo human cells, UMAP embeddings and clusters obtained from scVI and Scanpy respectively were utilized. Principal curves were obtained from TradeSeq^103^ using pseudotime and cellWeight values from slingshot. Gene expression and regulon activity were plotted as Z-Scores of pseudotime values from the fitGAM TradeSeq function using 6 knots.

### Ingenuity Pathway Analysis

Differentially expressed genes were fed into Ingenuity Pathway Analysis (Qiagen) to determine significant canonical, functional, and upstream signaling terms^123^. Activation Z-scores and p-values of curated significant (p < 0.05) terms were used for plotting with ggplot2^100^.

### Cell-Cell Interaction Analysis

For cell-cell interaction analysis between in vitro and in vivo hGPCs, the differential NicheNet pipeline was adapted ^38^. This analysis utilized human cells from the in vitro GPC stage and both human and mouse cells from the 19 week human glial chimeric corpus callosum. For the receiver cells (in vitro GPC4 and in vivo GPCs and cGPCs that cluster within GPC4), all human genes were used for analysis. For sender cells, only one to one orthologs were utilized as determined using biomaRt to access ensembl ^104^. Gene expression in a cell type was deemed as positive if a gene was detected in >10% of cells. Differential ligand expression was determined between in vitro and in vivo niches using both mouse and human cells. For this the Seurat FindMarkers function was used iteratively as implemented in NicheNet via calculate_niche_de using the Wilcoxon rank sum test. For receiver cells, differentially expressed genes were determined prior using MAST

We next used human cell type clusters from the in vitro GPC stage prior to transplantation (**Fig. 2**) as well as the mouse and human cells in the chimeric mouse CC, to predict differential ligand signaling affecting hGPCs between both niches using the NicheNet package ^38^. We focused on GPCs classified as GPC4 (**Fig. 2D**) as receiver cells since this population comprises the most specified hGPCs in vitro as well as hGPCs and hcGPCs in vivo (**Fig. 4A**). Since these hGPCs are only exposed to human sender cells in vitro, but both mouse and human sender cells in vivo, we first filtered the genes in these cells to only one to one orthologs. This approach permitted comparable ligand expression in a cross-species manner. Since receiver GPCs are solely human, we were able to utilize their entire transcriptomes for assessing expression of receptors and downstream ligand targets. Transcriptional differences in the receiver cells were obtained from the prior analysis (**Fig. 4B**) whereas differential expression of ligands across niches were the composite of all combinations of pairwise Wilcoxon rank sum tests. Receiver hGPCs were then scored for downstream activation of ligands from NicheNet’s database. Displayed ligands were curated based on a composite of higher in vivo ligand activity (**Fig. 5C**), higher cell type expression of the ligand in vivo (**Fig. 5D**), and sufficient receptor expression in in vivo hGPCs (>10% expressed) as well as, in many cases, higher receptor expression (**Fig. 5E**). Finally, we constructed a network of these select ligands and their predicted in vivo hGPC-enriched targets (**Fig. 5F**, ligand-target weights > 0.05) curated for hGPC receptors of these ligands (from **Fig. 5E**), salient glial genes (from **Fig. 4B**), and active GPC regulons (from **Fig. 4C**).

## Materials availability

Cells and reagents generated in this study will be made available on request. All requests for materials and reagents should be directed to the Lead Contact. For some cellular reagents and plasmids, a completed Material Transfer Agreement may also be required.

## Data availability

All gene expression data have been submitted to GEO, accession number GSE285158. Data were aligned to the GRCh38 or GRCm39 genomes and annotated using Ensembl v.106. Expression data are also available and may be further interrogated at <u>GlialExplorer.io</u>. SCENIC rankings were downloaded from https://resources.aertslab.org/cistarget/.

## Code availability

Complete R, Python, and Bash Analysis Scripts may be accessed at https://github.com/CTNGoldmanLab/Glial_Chimera_Maturation. Further requests for information, data and code should be directed to and will be fulfilled by the Lead Contact.

## Acknowledgments

This work was supported by the Adelson Medical Research Foundation, and NIH grants R01NS110776 and R01AG072298 to S.G, as well as by both Sana Biotechnology, Inc. and CNS2, Inc. We thank Lorenz Studer (Memorial Sloan-Kettering) for his generous provision of C27 cells, and Casey Payne for expert assistance with cell culture.

## Author contributions

JNM, BM, and DCM cultured all cells; BM collected all samples for CUT&Tag and scATAC-seq; XW, BM, and JNM analyzed all CUT&Tag data; SJS transplanted all chimeric mice; JNM and SJS dissociated all chimeric mice, JNM isolated all cells for transplant and scRNA-seq captures; CCL and NPTH optimized the chimeric aligner and NPTH ran the pySCENIC analysis; JNM conducted all other bioinformatic analyses; JNM and SAG designed the study and wrote the paper. All authors approved the final manuscript.

