## Supplemental Figures for "Charting the transition from in vitro gliogenesis to the in vivo maturation of transplanted human glial progenitor cells"

**6 Supplementary Figures**

**Supplementary Figure 1** *Related to Figure 1*

In vitro human glial progenitor cell differentiation is orchestrated epigenetically

**Supplementary Figure 2** *Related to Figure 2*

Heterogeneity of cultured human GPCs

**Supplementary Figure 3** *Related to Figure 2*

Changes in chromatin accessibility mediate hGPC heterogeneity

**Supplementary Figure 4** *Related to Figure 3*

Composition of in vivo human glial chimeric white matter

**Supplementary Figure 5** *Related to Figure 3*

Chromatin accessibility in vitro predicts differentiation bias in vivo

**Supplementary Figure 6** *Related to Figure 5*

Resident mouse brain cells drive the specification and differentiation of human GPCs in vivo

**7 Supplementary Data files**

**Supplementary Data 1** *Related to Figure 1*

The transcriptional and epigenetic landscape of hGPC differentiation in vitro

**Supplementary Data 2** *Related to Figure 2 and Supplemetnary Figure 3*

hGPC subpopulations in vitro: gene and chromatin accessibility enrichment

**Supplementary Data 3** *Related to Figure 3*

Transcriptional characterization of hGPCs in vivo

**Supplementary Data 4** *Related to Figure 4 and Supplementary Figure 3*

In vivo vs in vitro GPC4 differential expression, regulon activity, IPA terms, and consensus with scATAC-Seq

**Supplementary Data 5** *Related to Figure 5*

Mouse cell type markers and differential in vivo vs in vitro GPC4 NicheNet cell-cell communication

**Supplementary Data 6** *Related to Figure 6*

In vivo oligodendrocyte differentiation, differential expression and regulon activity

**Supplementary Data 7** *Related to Figure 6*

In vivo astrocyte differentiation, differential expression and regulon activity

**Supplementary Figure 1** *Related to Figure 1*

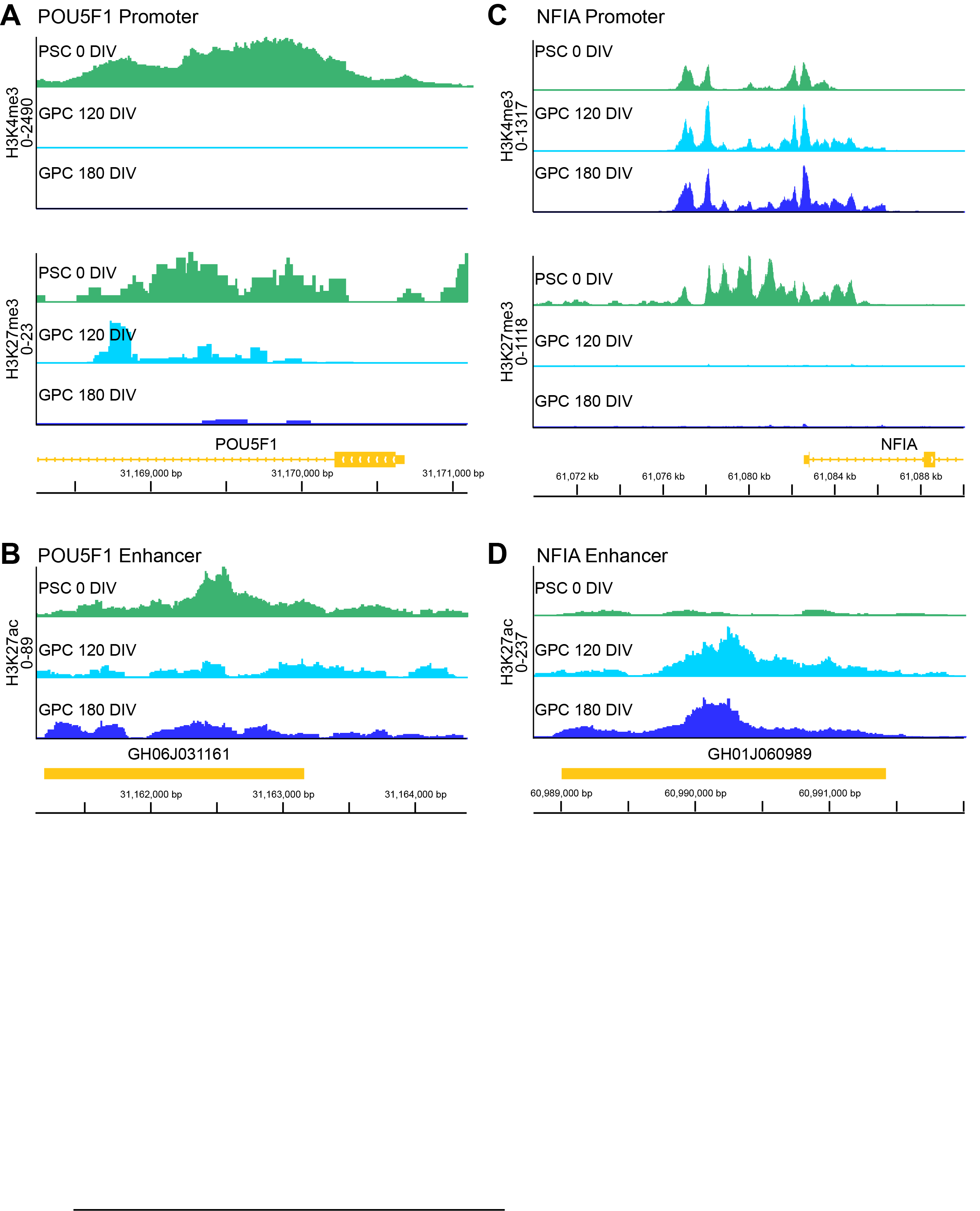

**In vitro human glial progenitor cell differentiation is orchestrated epigenetically**

Cleavage under targets & tagmentation (CUT&Tag) of PSC, 120 DIV GPCs, and 180 DIV GPCs for H3K4me3, H3K27me3, and H3K27ac. **A.** H3K4me3 and K3K27me3 normalized coverage plots at the POU5F1 promoter. **B.** Normalized H3K27ac coverage at POU5F1 enhancer GH06J031161. **C.** H3K4me3 and K3K27me3 normalized coverage plots at the NFIA promoter. **D.** Normalized H3K27ac coverage at NFIA enhancer GH01J060989.

**Supplementary Figure 2** *Related to Figure 2*

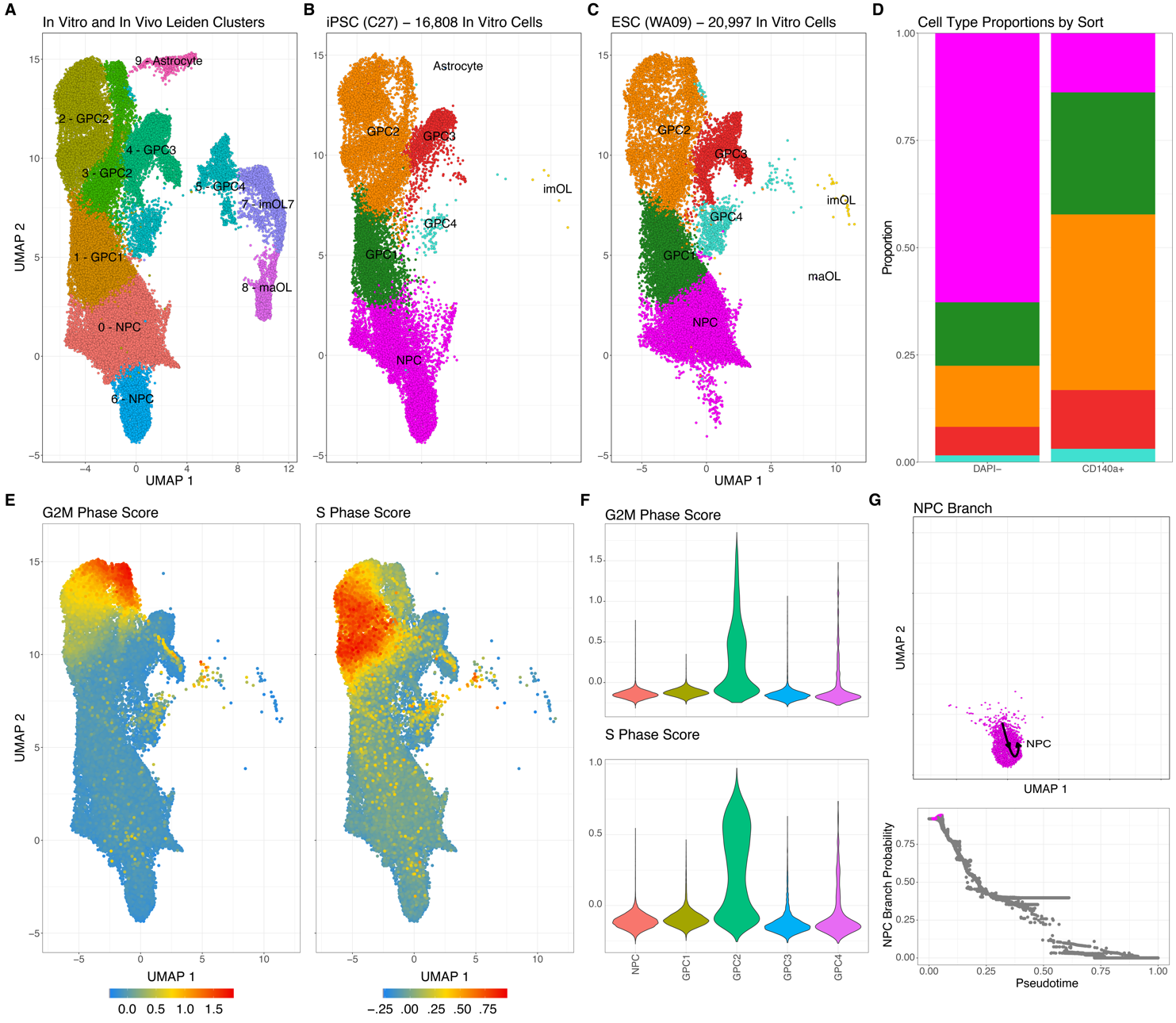

**Heterogeneity of cultured human GPCs**

**A.** Leiden clusters and manually assigned subpopulation names of cultured GPCs. **B-C.** Subpopulations of cultured iPSC- **(B)** and **(C)** ESC-derived GPCs. **D.** Proportions of FACS DAPI depleted or both DAPI depleted and CD140a+ cultured GPCs. **E-F.** G2M and S phase scores calculated in Seurat plotted as feature **(E)** or violin plot **(F)**. **G.** UMAP with overlaid trajectories (top) and branch probability plots across pseudotime of NPC lineage calculated with Palantir. *GPCs: Glial progenitor cells, NPCs: Neural progenitor cells, imOLs and maOLs: Immature and mature oligodendrocytes.*

**Supplementary Figure 3** *Related to Figure 2*

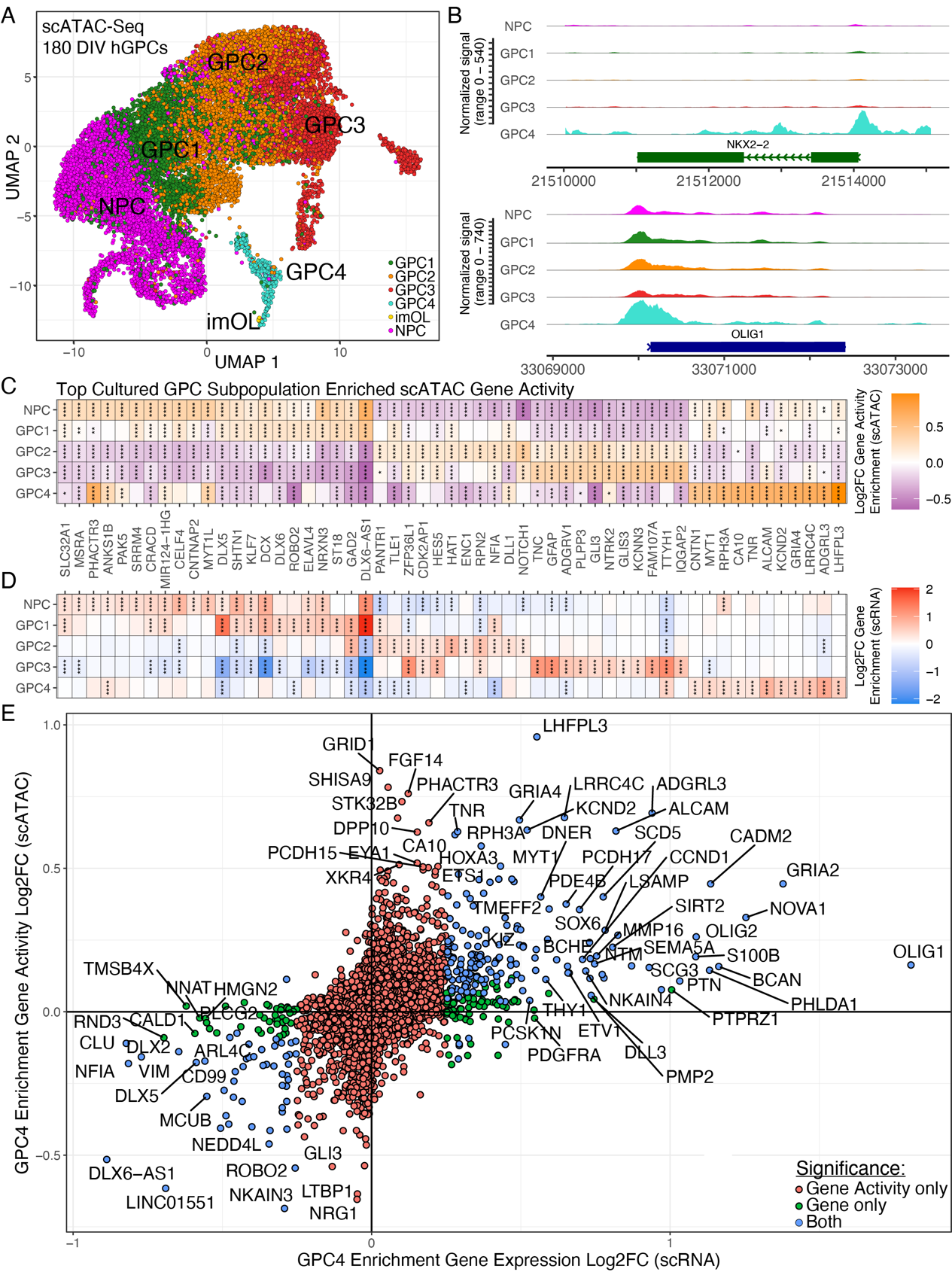

**Changes in chromatin accessibility mediate hGPC heterogeneity**

**A.** scATAC-seq UMAP of 180 DIV hESC-derived (WA09) hGPCs with label transferred cell populations (n=1). **B.** Coverage plots of GPC4 accessible genes. **C-D.** Heatmaps of the top enriched gene activities **(C)** calculated from scATAC-seq for each subpopulation aligned with gene expression enrichment **(D)** from cultured hGPC scRNA-seq. **E.** Scatter plot of GPC4 differential gene expression enrichment vs GPC4 differential chromatin accessibility. Blue dots are significant for both assays, red are significant only for gene activity (scATAC), and green only for gene expression (scRNA). *GPCs: Glial progenitor cells, NPCs: Neural progenitor cells, imOLs and maOLs: Immature and mature oligodendrocytes.* FDR: ****<0.00001, ***<0.0001, **<0.001, *<0.01.

**Supplementary Figure 4** *Related to Figure 3*

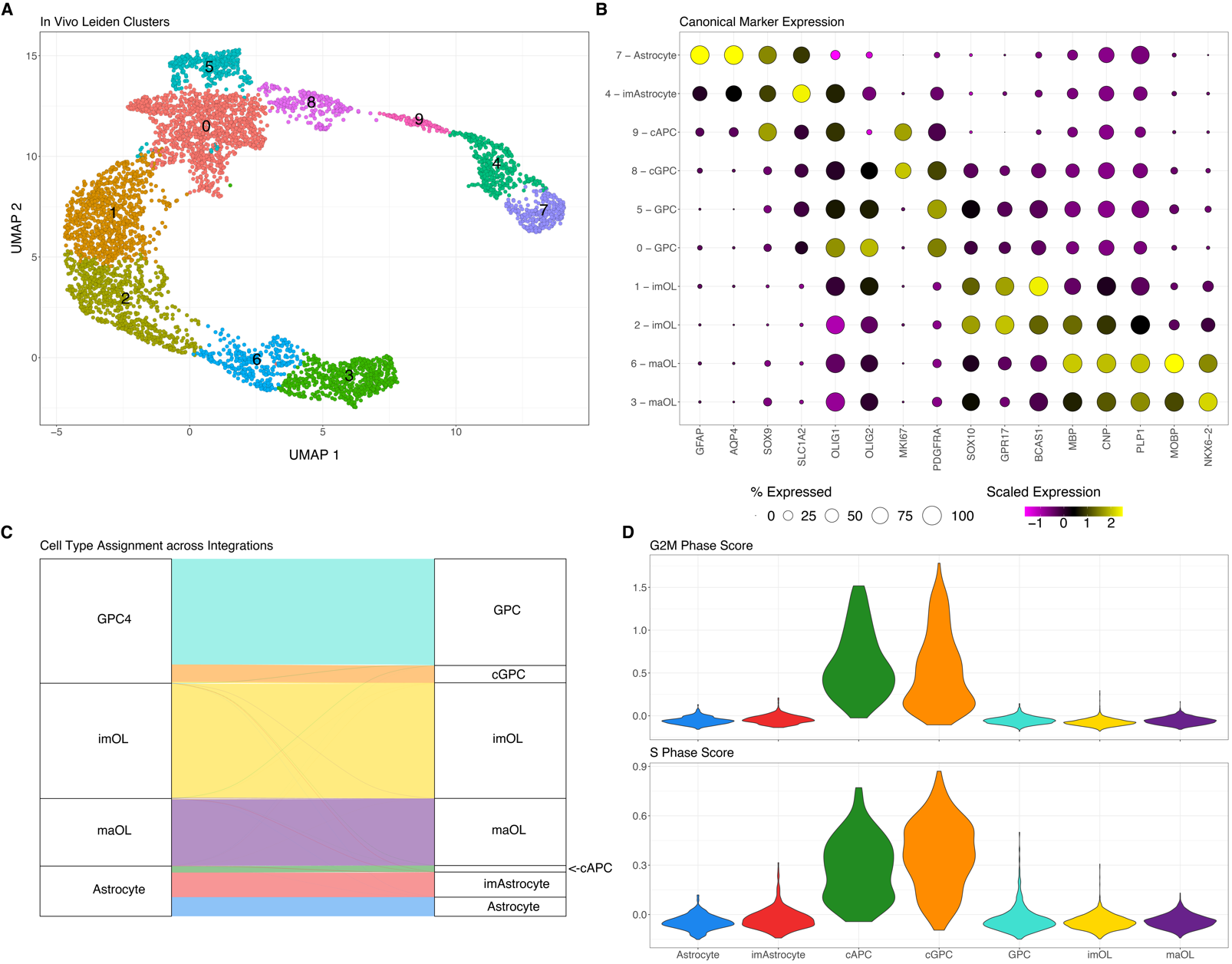

**Composition of in vivo human glial chimeric white matter**

**A.** Leiden clusters and assigned cell types of human cells extracted back from 19wk old shiverer corpus callosum. **B.** Cell type markers of human leiden clusters and their manually assigned aggregate cell types. **C.** Alluvial plot showing cell type calling between combined (**Fig. 2**) and in vivo-alone integrations (**Fig. 3**). **D.** G2M and S phase scores calculated in Seurat. *GPCs: Glial progenitor cells, imOLs and maOLs: Immature and mature oligodendrocytes, cGPC: Cycling GPCs, APC: Astrocyte progenitor cells, cAPC: Cycling APCs, imAstrocyte: Immature astrocytes.*

**Supplementary Figure 5** *Related to Figure 3*

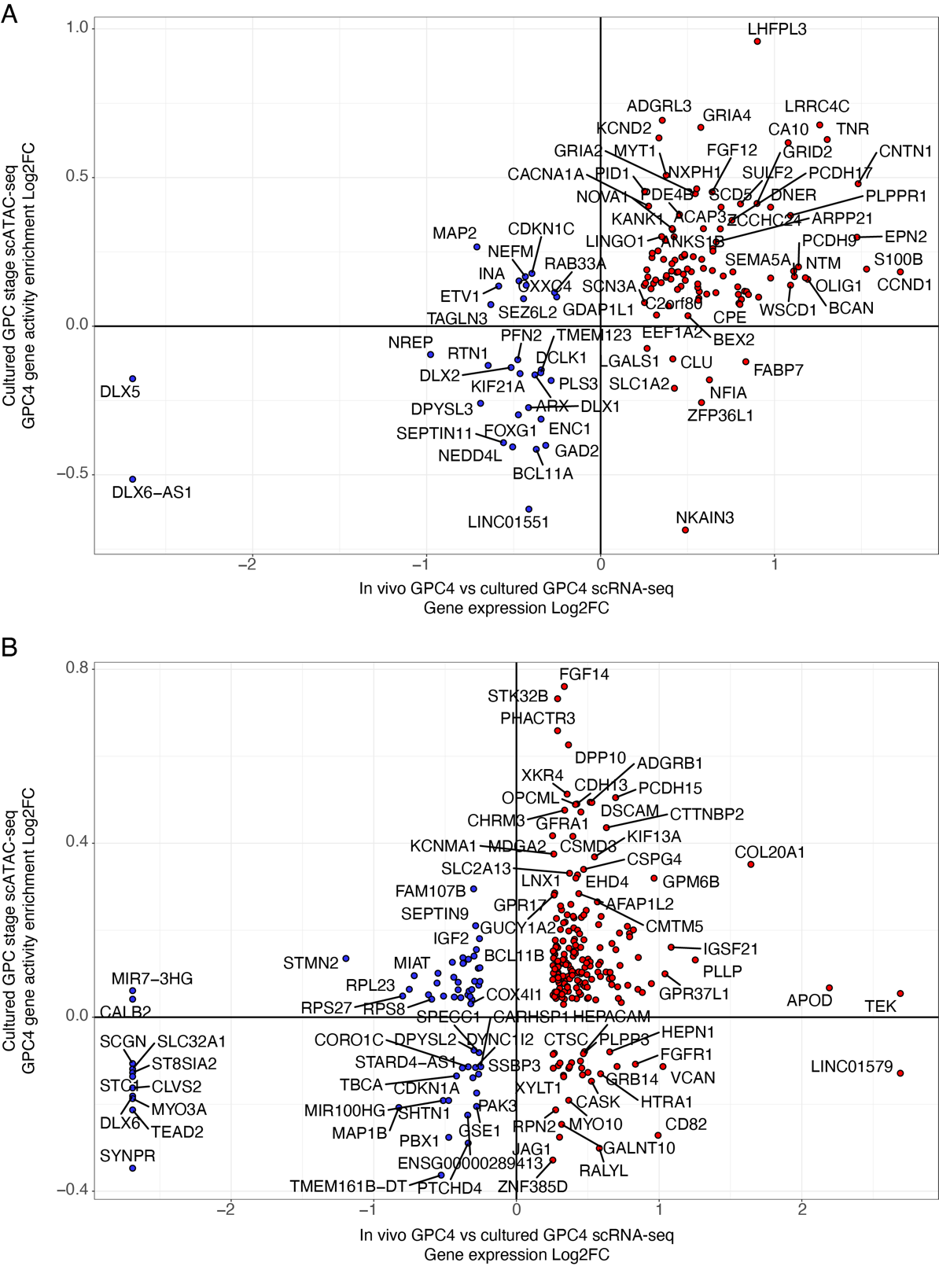

**Chromatin accessibility in vitro predicts differentiation bias in vivo**

            Scatter plots of in vitro GPC4 stage scATAC-seq gene activities, plotted as log_2_FC enrichment (GPC4 vs remaining in vitro populations, from **Supplementary Fig. 3E**) on the Y-axes, relative to in vivo vs in vitro differential gene expression log_2_FCs (from **Fig. 4B**) on the X-axes. **A.** The plotted genes are significant for differential expression between in vivo and in vitro GPC4s and were also significant for their relative gene activities and differential enrichment in cultured GPC4s, compared to other cultured GPC subpopulations. A selective preponderance of oligodendroglial lineage genes is apparent. **B.** Genes plotted are significant for differential expression between in vivo and in vitro GPC4s, and were also significant for their differential gene activities of GPC4 stage cells relative to other GPCs in vitro, but *not* for enrichment of corresponding RNA expression in cultured GPC4s. This suggests the availability of maturation-associated genes poised for activation-induced expression in response to tissue-derived cues after in vivo transplantation.

**Supplementary Figure 6** *Related to Figure 5*

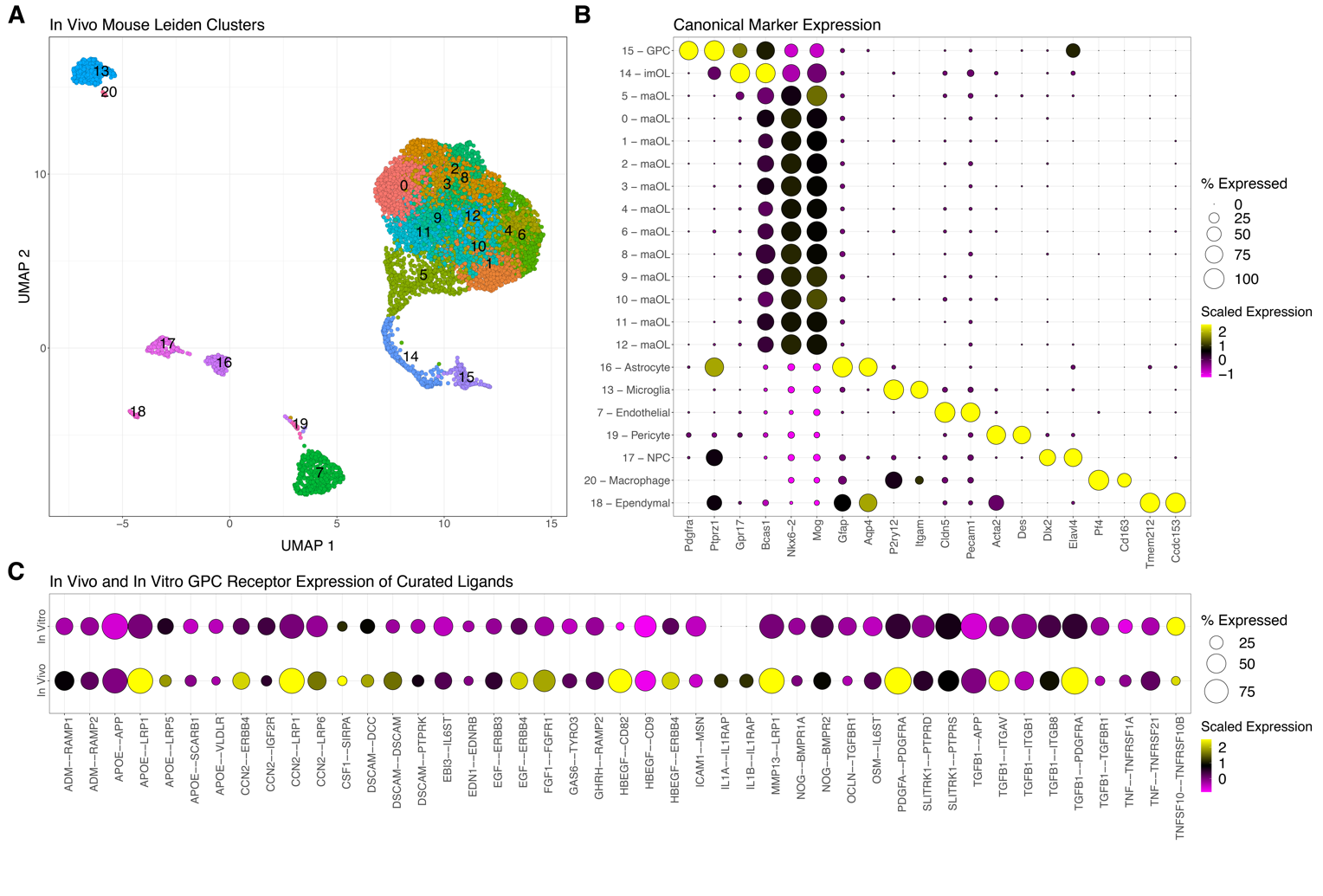

**Resident mouse brain cells drive the specification and differentiation of human GPCs in vivo**

**A.** Leiden clusters of integrated mouse cells in engrafted shiverer corpus callosum. **B.** Cell type markers of mouse leiden clusters and their manually assigned aggregate cell types. **C.** In vitro and in vivo GPC4 scaled and percent expression of receptors associated with identified active in vivo ligands.
